# Targeting the water network in cyclin G associated kinase (GAK) with 4-anilino-quin(az)oline inhibitors

**DOI:** 10.1101/2020.03.06.976563

**Authors:** Christopher R. M. Asquith, Graham J. Tizzard, James M. Bennett, Carrow I. Wells, Jonathan M. Elkins, Timothy M. Willson, Antti Poso, Tuomo Laitinen

## Abstract

Water networks within kinase inhibitor design and more widely within drug discovery are generally poorly understood. The successful targeting of these networks prospectively has great promise for all facets of inhibitor design, including potency and selectivity on target. Here we describe the design and testing of a targeted library of 4-anilinoquinolines for use as inhibitors of cyclin G associated kinase (GAK). The GAK cellular target engagement assays, ATP binding site modelling and extensive water mapping provide a clear route to access potent inhibitors for GAK and beyond.

---

Inhibitor specificity and potency both present significant challenges in the development of kinase inhibitors and exploitation of kinase targets as potential therapies. The target profiles of commercial broad-spectrum kinase inhibitors are often challenging to rationalize in terms of direct molecular activity profiles. The large number of protein kinases present in the kinome (>500) makes this task even more difficult.^[1–2]^ Protein kinases that are functionally distinct yet retain key structural features can show overlapping ligand preferences, both with near and distant neighbours on the kinome tree.^[3]^

A number of strategies have been explored to address the specificity problem in kinases.^[4]^ These approaches include exploitation of size of the kinase gatekeeper residue, the disposition of the DFG-loop, chemotype selectivity, non-covalent interactions, salt-bridge, solvation, etc. Another method employed to obtain selective kinase inhibitors has been to target an allosteric site rather than the conserved ATP-binding site that is traditionally targeted and one that is unique to the kinase in question. Additionally, synthesizing inhibitors capable of covalent binding to a proximal cysteine has been successful to achieve selectivity over close family members. While each approach offers a possibility to achieve selectivity there is still a significant overall challenge to reach a successful endpoint. Identification and design of allosteric binders is challenging and most are found serendipitously.^[5]^ While inhibitors that modify the kinase covalently, first there has to be a residue capable of modification and then a way to assess the biological impact of that permanent modification.^[6]^

There has been a growing appreciation that solvent specific effects can make significant contributions to ligand binding. The impact of these effects are however, usually retrospectively applied as a post-rationalization of the observed results.^[7–9]^ Development of software to model these effects has progressed with enveloping distribution sampling (EDS), free energy perturbation (FEP) theory and displaced-solvent functional (DSF) all recently being developed.^[10–11]^ However, the use of WaterMap by Schrödinger, Inc. has significantly advanced the ability to study direct solvation effects such as solvent replacement.^[12]^

WaterMap couples molecular dynamics simulations with statistical thermodynamic analysis of water molecules within a protein structure. The method can provide insight not possible with traditional docking approaches. WaterMap was first used to rationalize the structure activity relationships of triazolylpurines binding to the A2A Receptor.^[13]^ Specifically, *n*-butyl and *n*-pentyl substituents resulted in unanticipated potency gains, but these structure activity relationships could not be readily explained by ligand−receptor interactions, steric effects, or differences in ligand desolvation.

WaterMap has also been utilized in a number of kinase inhibitor programs including Src family kinases, Abl/c-Kit, Syk/ZAP-70, and CDK2/4 in order to rationalize kinase selectivity.^[13]^ Water modifications have also been shown to have a key effect in pivotal EGFR mutations showing resistance to afatinib and erlotinib.^[14]^ Molecular modelling studies on a PDGF-Rβ homology model using prediction of water thermodynamics suggested an optimization strategy for the 3,5-diaryl-pyrazin-2-ones as DFG-in binders by using a phenolic hydroxyl group to replace a structural water molecule in the ATP binding site.^[15]^

This water network principle has also been applied to bosutinib, as the 3-cyanoquinoline allows the compound to engage with a pair of conserved structured water molecules in the active site of Src.^[16]^ This post-rationalization approach hints at a more powerful use of this idea in the design of new inhibitors to exploit the water network hypothesis in its entirety to address both potency and selectivity.^[13]^

In our work developing a selective chemical probe for Cyclin G associated kinase (GAK), a member of the NAK sub-family.^[17–21]^ We observed two matched pairs of experimental results that did not fit with the expected structure activity relationship observed for GAK and the wider 4-anilinoquinoline scaffold (**Figure 1**).^[17]^

**Figure 1.**
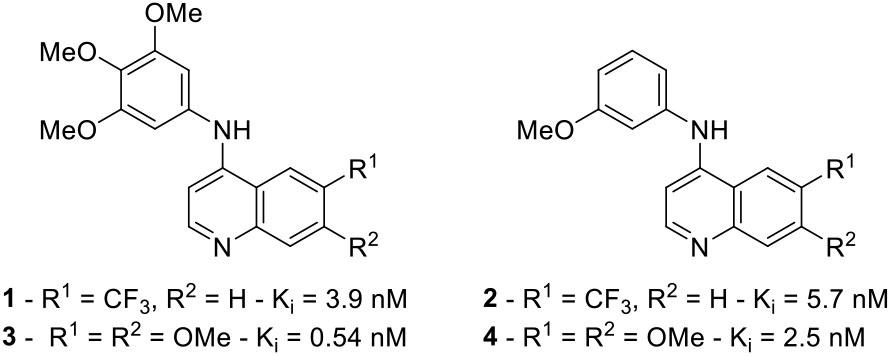
4-Anilinoquinoline GAK inhibitor matched pairs with GAK activity shown.

GAK is involved in a plethora of biological processes including cell cycle progression,^[22]^ Parkinson’s disease,^[23]^ osteosarcoma^[24]^ and prostate cancer.^[18,25]^ Although GAK is a historically under-studied kinase,^[26]^ recent probe development efforts identified potent and selective inhibitors of GAK from multiple chemotypes, including *iso*thiazolo[5,4-*b*]pyridines,^[27–28]^ 4-anilinoquinolines and 4-anilinoquinazolines.^[17–20]^ These chemical probes may be used in the future to study of the biology of GAK.^[17,29]^ In contrast to probes, several FDA-approved drugs (**Figure 2**), including EGFR inhibitors gefitinib and erlotinib, have been reported to show off-target GAK activity in the low nanomolar range (**Figure 2**).^[30]^

**Figure 2.**
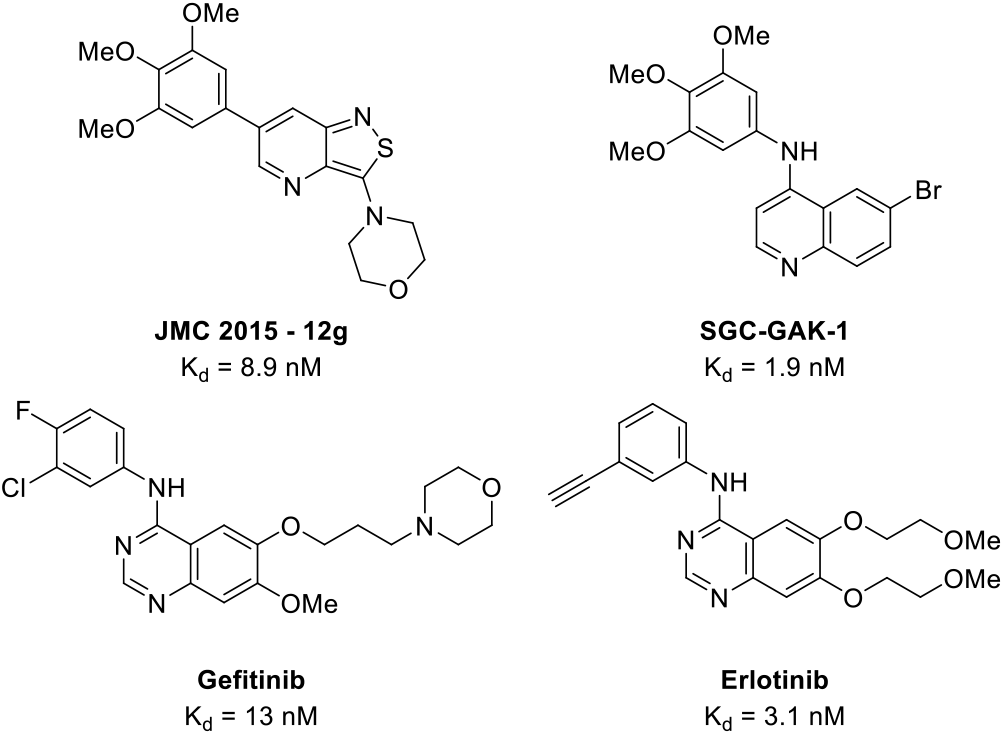
Previously reported GAK inhibitors.

## Results

The differently substituted matched pairs of trimethoxy- and *meta*-methoxy-substituted quinolines **1**-**4** showed potent binding to GAK with good NAK-family selectivity despite significant differences in both steric and electronic footprints.^[17]^ We looked for an alternative explanation for our observed equipotency and found through the application of WaterMap that there was a high-energy hydration site in the *p*-loop of the active site which is enabling both enhanced potency and NAK-family selectivity.

To further investigate this, we first looked at each NAK family member and observed that all have distinct water networks (**Figure 3A-D**). AAK1 and BIKE have a high degree of sequence identity in the kinase domain.^[21]^ There are some key residue differences located at the hinge region of each NAK family member: Leu125 (GAK) / Phe100 (STK16) / Tyr132 (BIKE) / Phe128 (AAK1), and Phe101 in STK16 in place of Cys126 (GAK)/ Cys133 (BIKE) / Cys126 (AAK1). There are also alterations in the back pocket, where polar Thr123 and other bulkier residues like Leu98 (STK16) are located in case of GAK and Met130 and Met126 in case of BIKE and AAK1. Respectively, these alterations have significant impacts on the binding modes of inhibitors (**Figure 3A-B**).^[20]^ While the other three family members have lipophilic residues that have no ability to behave as a hydrogen bond donor/acceptors, GAK has Thr123 that can perform both functions. Moreover, the rotamer of Thr123 is likely fixed in place by an internal hydrogen bond to the backbone carbonyl group of Glu124 (**Figure 3C-D**).

**Figure 3.**
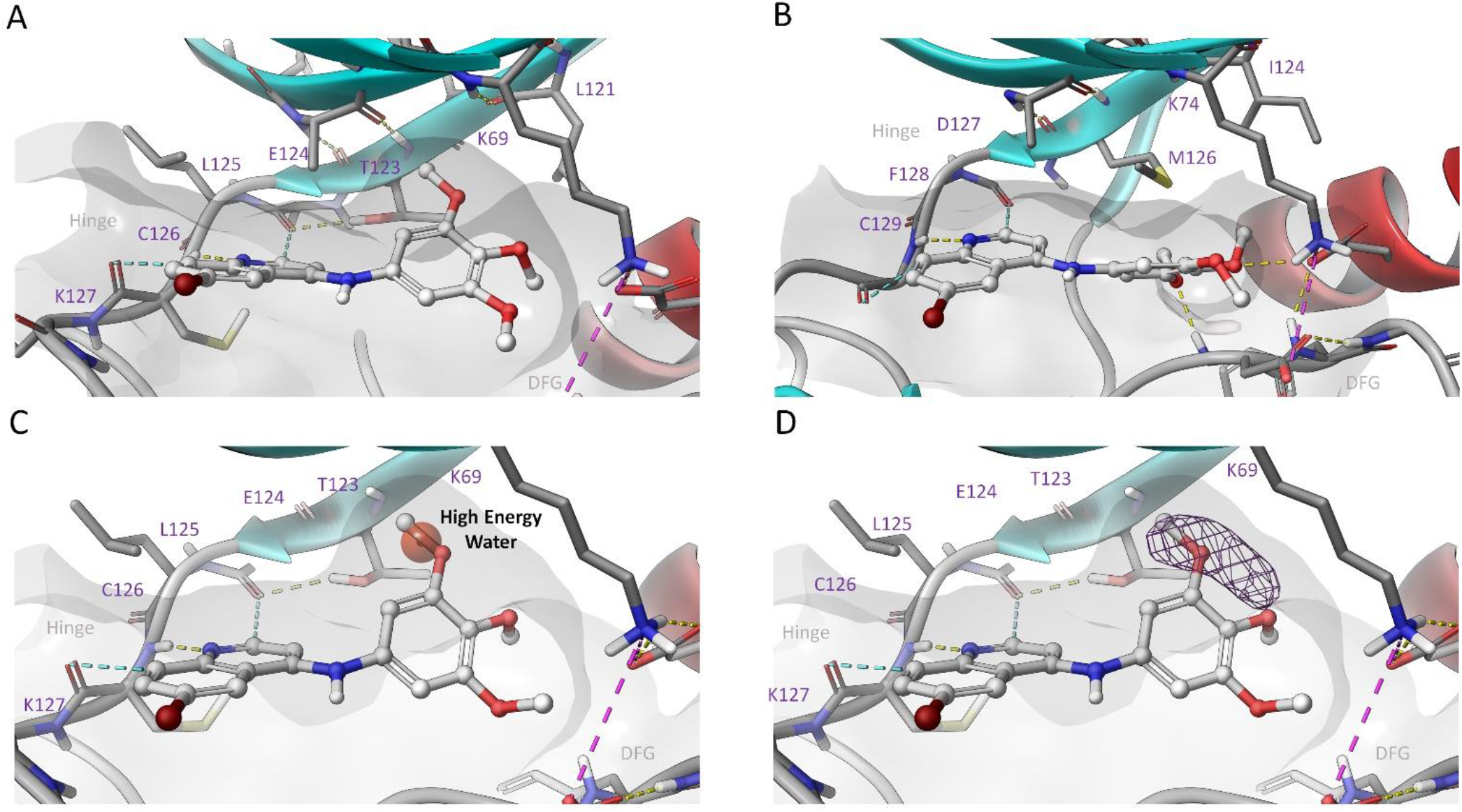
**A**: shows additional space close to beta-sheet at the back pocket occupied in favourable docking pose of compound SGC-GAK-1. **B**: shows that in case of AAK1 the corresponding back pocket area is filled by methionine side chain (same location as Thr123 in GAK) for comparison. **C**: red sphere shows location of high energy (5.9 kcal/mol) hydration site in GAK. **D**: shows that high energy hydration site is actually dewetted (very low water occupancy) in WaterMap simulation.

This also gives rise to differences in the predicted water networks of the four NAK kinases (**Figure 4A-D**). However, only GAK has a high energy hydration site likely due to small pit created by the interaction between Thr123 and Glu124 (**Figure 4A**). A hydration site is considered to be high in energy when the relative free energy value from WaterMap analysis has a high positive value i.e. 5.9 kcal/mol in case of GAK. There can be two cases when the hydration site has unfavourable high energy. First, water solvating hydrophobic enclose such as in the case of GAK third strand of beta sheet (β3) between Lys69 and Thr123. This type of cavity is energetically unfavourable (due to enthalpy) because water molecules cannot form a full complement of hydrogen bonds with surroundings. Second, is where the binding is entropically unfavourable, partly due to the closed nature of the hydration site. This is where water can form hydrogen bonds but suffers from reduced number of hydrogen bond configurations with protein groups and partner waters. Replacement of a water from such hydrations sites often may lead to improvement in affinity. Despite 17 sequence differences between GAK and AAK1/BIKE in the ATP binding domain;^[21]^ the only significant difference in the water network is the presence of the high-energy water in the *p*-loop region (**Figure 3A-D**). The most structurally diverse NAK family member STK16 has 64 differences in the kinase sequence compared to the other NAK kinases in the ATP binding domain. STK16 has a significantly different water network compared to the other NAK family members but does possess a high energy water that is in a 7 Å out of plane orientation related to the GAK protein structure (**Figure 4A** & **4D**).

**Figure 4.**
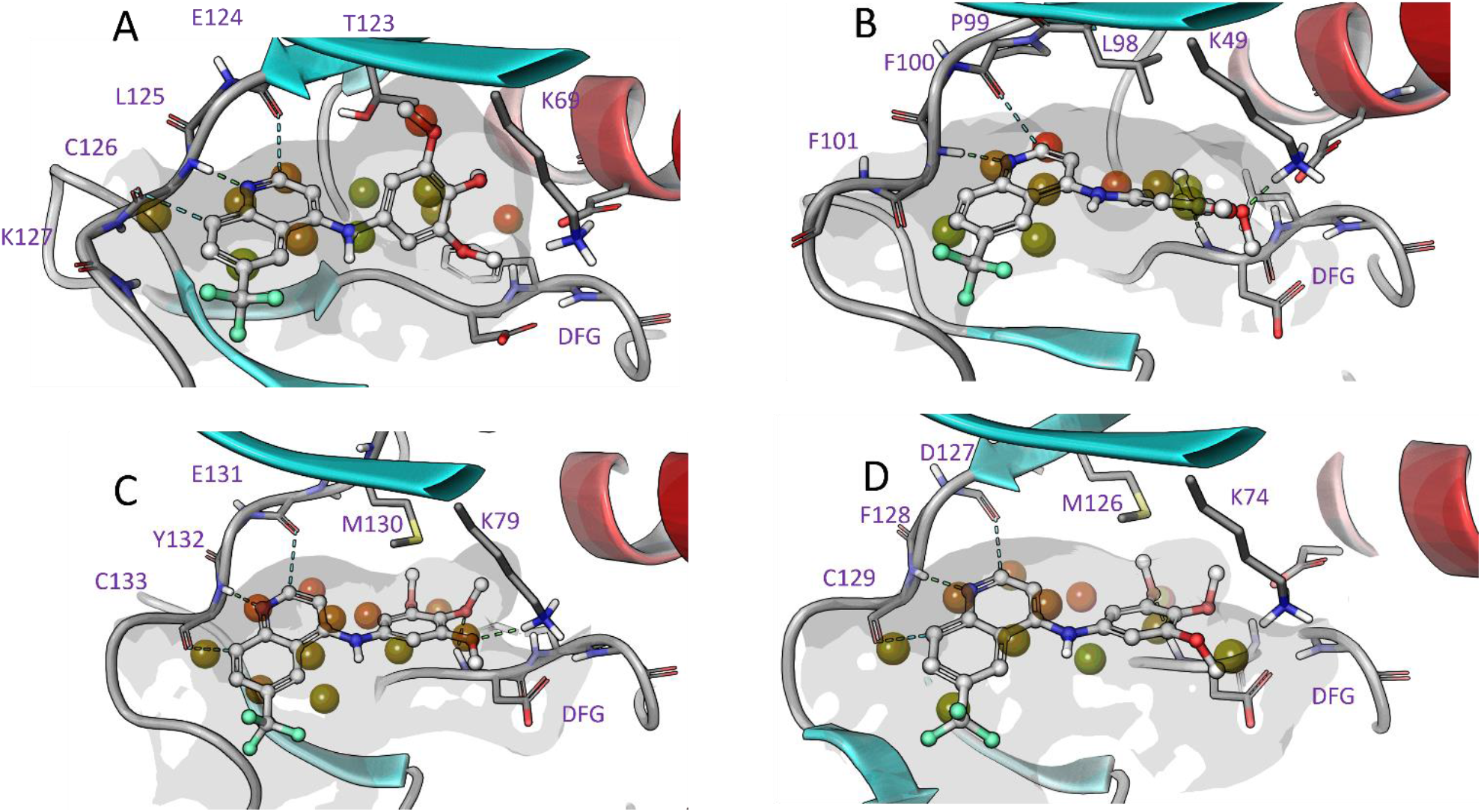
Molecular Modelling **1** in the water network of NAK family members using WaterMap (high energy waters are highlighted in red, lower energy waters are in green): **A**: GAK (PDB:5Y80); **B**: STK16 (PDB:2BUJ); **C**: BIKE (PDB: 4W9X); **D**: AAK1 (PDB: 5TE0)

Through a series of prospective simulations using WaterMap we identified that this additional high energy hydration site next to the spine of the ATP binding pocket makes the shape of the GAK binding back pocket rather different to other NAK kinases studied (**Figure 4**).^[31]^ A hydrophobic furrow is formed next to Thr123, where the sidechain is hydrogen bonded to the backbone carbonyl group of Glu124, and is rather small in size overall. While in case of other NAK kinases there is either a bulky methionine (AAK1 and BIKE) or Ile110 in the case of STK16 (**Figure 3A-B**). As a consequence, in the case of GAK, this hydrophobic site can be readily occupied by a ligand substituent without side chain rearrangements.

The water network simulation of the 6-(trifluoromethyl)-*N*-(3,4,5-trimethoxyphenyl)quinolin-4-amine (**1**) vs *meta*-methoxy derivative (**2**) demonstrated plausibility of the high energy water displacement to explain the almost equipotent high GAK activity (**Figure 5A-B**). The slight increase in potency could be attributed to the rearrangement of the catalytic lysine to sit between the two additional methoxy groups in **1**. Reforming of the water contacts by removal of the methyl group from the methoxy group (**5**) showed an encouraging ability to lock the phenolic −OH in place in the preferred docking pose (**Figure 5C**). Compound **5** could also be forced to adopt a higher energy conformation where the −OH interacts with the catalytic lysine (**Figure 5D**). The new H-bond lysine interaction is not compensating the two H-bonds replacing the high energy water. Although metabolically liability of a phenolic −OH is well known, it is a strong proof of concept substitution.^[32]^

**Figure 5.**
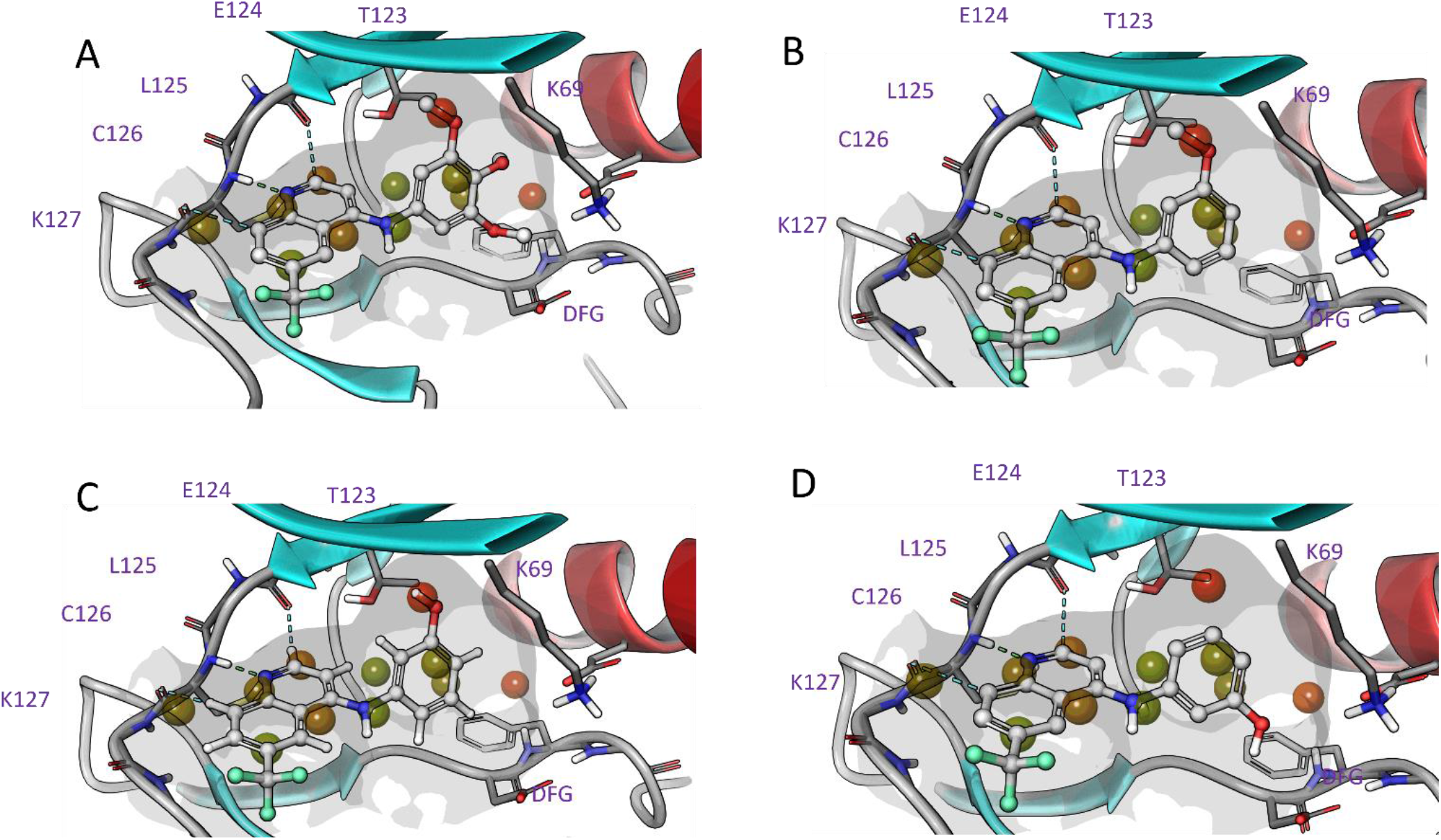
WaterMap modelling of GAK where high energy waters are highlighted in red, lower energy waters are in green (PDB:5Y80), with: A: **1**; B: **4**; C: **5**; D: **5**

To further probe the structural requirements for GAK activity and the water network, we synthesized a series of analogs (**Scheme 1**). 4-Anilinoquinolines were prepared by heating the corresponding 4-chloroquinoline derivative and substituted aniline in ethanol with Hünigs base and refluxed overnight.^[17–20,33–34]^ The synthesis afforded products **1**-**34** in good yield (58-86%). Interestingly, the more activated reaction to 6-bromo-4-chloroquinoline-3-carbonitrile was completed in two hours to afford **31** in good yield (69%). The hydrogenations of the nitro derivatives **7** and **21** to form **8** and **22** were routine with good to excellent yields (69 and 94% respectively).

**Scheme 1.**
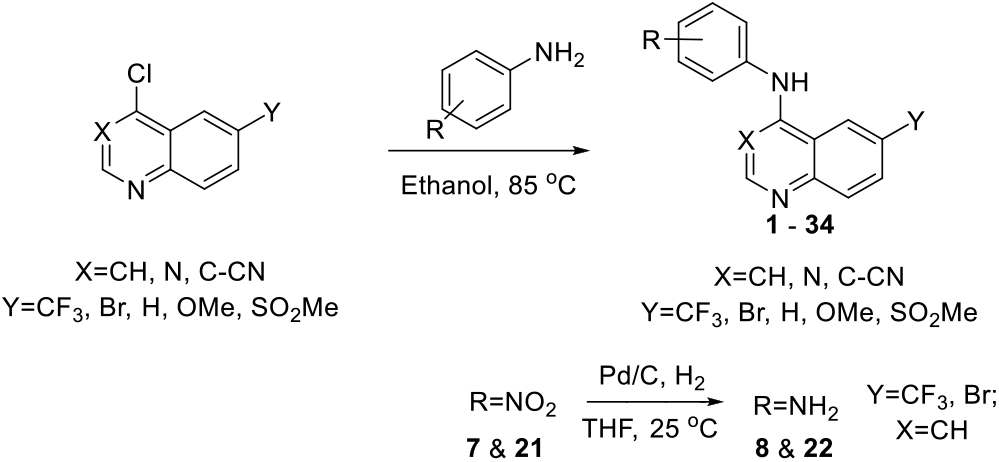
General synthetic route to analogs **1** − **34**.

The compounds were initially screened for activity on the kinase domains of all four members of the NAK family members (GAK, AAK1, BMP2K, and STK16) using a time-resolved fluorescence energy transfer (TR-FRET) binding displacement assay in a 16-point dose response format to determine the inhibition constant (K_i_). The compounds were then screened in a cellular target engagement assay nanoluciferase (nanoLuc) bioluminescence resonance energy transfer (BRET) (NanoBRET) assay for GAK.^[17–18,35–36]^ The sensitive nature of the assay allows for a complimentary comparison to the TR-FRET to observesubtle trends in the GAK inhibition profile.

The background provided by **1**-**4** and **Figure 3A-D** led us to probe around the 6-trifluorometyl substituted quinolines with some small modifications looking for potential hydrogen bond interactions to further boost the initial potency in the structure activity relationships observed (**Figure 1**).^[17]^ The trimethoxy compound **1** demonstrated high affinity for GAK and good selectivity against the rest of the NAK family members. A switch from the trimethoxy-(**1**) to the *meta*-methoxy-substitution (**2**) showed almost equipotency consistent with previous reports.^[17]^

The GAK NanoBRET showed an almost 3-fold drop in cellular target engagement from **1** to **2**. This effect is still impressive as the *meta*-methoxy derivative **2** is able to substitute the trimethoxy functionality despite not occupying the same space orientation.^[16]^ A switch from *meta*-methoxy (**2**) to *meta*-hydroxy (**5**) showed a slight increase in GAK binding but not as significantly as the modelling predicted (**Table 1**).

**Table 1.** Investigation of matched pairs of *meta*-substituted quinolines.

| Cmpd | R<sup>1</sup> | R<sup>2</sup> | R<sup>3</sup> | R<sup>4</sup> | R<sup>5</sup> | Binding displacement assay K<sub>i</sub> (μM)<sup>b</sup> |  |  |  | NAK selectivity<sup>c</sup> | GAK NanoBRET IC<sub>50</sub> (μM)<sup>d</sup> |
| --- | --- | --- | --- | --- | --- | --- | --- | --- | --- | --- | --- |
|  |  |  |  |  |  | GAK | AAK1 | BMP2K | STK16 |  |  |
| **1<sup>a</sup>** | OMe | OMe | OMe | H | CF<sub>3</sub> | 0.0039 | 54 | >100 | 17 | 4300 | 0.17 |
| **2<sup>a</sup>** | OMe | H | H | H | CF<sub>3</sub> | 0.0057 | 14 | 23 | 18 | 2500 | 0.48 |
| **5** | OH | H | H | H | CF<sub>3</sub> | 0.0047 | 2.6 | 5.8 | 3 | 550 | 0.23 |
| **6** | NO<sub>2</sub> | H | H | H | CF<sub>3</sub> | 0.21 | >100 | >100 | >100 | 480 | >5 |
| **7** | NH<sub>2</sub> | H | H | H | CF<sub>3</sub> | 0.12 | 6.8 | 17 | 45 | 57 | 2.7 |
| **8** | NMe<sub>2</sub> | H | H | H | CF<sub>3</sub> | 0.65 | >100 | >100 | 13 | 20 | >5 |
| **9** | NO<sub>2</sub> | H | H | H | H | 0.17 | 38 | >100 | >100 | 220 | >5 |
| **10** | NMe<sub>2</sub> | H | H | H | H | 0.97 | 29 | >100 | >100 | 30 | >5 |
| **11** | CH<sub>2</sub>OH | H | H | H | CF<sub>3</sub> | 0.06 | 10 | 15 | 20 | 170 | 1.7 |
| **12** | H | H | H | CH<sub>2</sub>OH | CF<sub>3</sub> | 0.041 | 8.5 | 16 | 5 | 120 | 2.3 |

<sup>a</sup>Consistent with previous report<sup>[17]</sup>; <sup>b</sup>TR-FRET-based ligand binding displacement assay; <sup>c</sup>(NAK/GAK); <sup>d</sup>IC<sub>50</sub> (μM) (n = 2, mean average)

We then switched to a *meta*-nitro group (**6**) or *meta*-amino group (**7**) to investigate the possibility of forming multiple hydrogen bonds.^[37]^ However, both **6** and **7** proved to be relatively ineffective with a 45 and 25-fold decrease in GAK binding respectively and a low micromolar cellular target engagement. The addition of two methyl groups to the amine (**8**) demonstrated a 5-fold decrease in GAK potency (K_i_ = 650 nM). The removal of the trifluoromethyl group of **7** and **8** to form **9** and **10** respectively showed the same trend in GAK potency but a 4-fold increase in selectivity (**8** vs **10**). We returned to the hydroxy moiety and inserted a methyl linker between the aniline with the aim to effectively generate a hydrogen bond interaction with Thr123. The methanol analogs **11** and **12** proved to be 10-fold weaker on GAK, likely due to spatial constraints in the top pocket of the ATP binding site.

The NAK family selectivity pattern observed with **1**-**5** hinted at a potential water displacement. With this in mind we explored a series of *meta*-methoxy and *meta*-hydroxy matched pairs (**Table 1**). Compound **2** has a favourable NAK family profile towards GAK and has previously been shown to have a narrow spectrum profile across the kinome.^[19]^ The 6-bromoquinoline (**13**-**14**) is the optimal substituent for GAK inhibition on this scaffold (**Table 2**).^[17–19,34]^ This substitution offers optimal GAK binding in the solvent exposed mouth of the ATP binding site and potent GAK inhibition at lower concentrations. The switch from the trimethoxy to the *meta*-methoxy substitution (**13**) was equipotent to the binding of **2** on GAK; but with a corresponding drop off in NAK selectivity and NanoBRET activity. The switch from **13** to the hydroxy substituted compound **14** shows an almost 3-fold increase in K_i_ which was more pronounced in the cellular assay with an almost 10-fold increase from an IC_50_ of 218 nM to 26 nM.

**Table 2.** Comparative results based on compound **1** and **2**.

| Cmpd | R<sup>1</sup> | R<sup>2</sup> | Binding displacement assay K<sub>i</sub> (μM)<sup>b</sup> |  |  |  | NAK selectivity<sup>c</sup> | GAK NanoBRET IC<sub>50</sub> (μM)<sup>d</sup> |
| --- | --- | --- | --- | --- | --- | --- | --- | --- |
|  |  |  | GAK | AAK1 | BMP2K | STK16 |  |  |
| **2<sup>a</sup>** | OMe | CF<sub>3</sub> | 0.0057 | 14 | 23 | 18 | 2500 | 0.17 |
| **5** | OH | CF<sub>3</sub> | 0.0047 | 2.6 | 5.8 | 3.0 | 550 | 0.48 |
| **13** | OMe | Br | 0.0029 | 3.0 | 7.0 | 32 | 1000 | 0.22 |
| **14** | OH | Br | 0.0012 | 0.50 | 2.0 | 5.7 | 420 | 0.026 |
| **15** | OMe | H | 0.027 | 25 | >100 | >100 | 930 | 0.60 |
| **16** | OH | H | 0.012 | 6.6 | 30 | >100 | 550 | 0.44 |
| **17** | OMe | OMe | 0.0079 | 9.1 | 26 | >100 | 1300 | 0.17 |
| **18** | OH | OMe | 0.0057 | 1.6 | 7.3 | 27 | 280 | 0.11 |
| **19** | OMe | SO<sub>2</sub>Me | 0.030 | 23 | 19 | >100 | 630 | 1.0 |
| **20** | OH | SO<sub>2</sub>Me | 0.023 | 5.1 | 3.5 | 1.2 | 520 | 1.0 |

<sup>a</sup>Consistent with previous report<sup>[17]</sup>; <sup>b</sup>TR-FRET-based ligand binding displacement assay; <sup>c</sup>(NAK/GAK); <sup>d</sup>IC<sub>50</sub> (μM) (n = 2, mean average)

The removal of the 6-bromo to the unsubstituted quinoline analog **15** resulted a in 10-fold drop in potency of *meta*-methoxy (**15**); with the hydroxy (**16**) only giving a limited increase in GAK affinity. The NanoBRET results for **15** and **16** were weaker but consistent with the trend observed. The 6-methoxy-*N*-(3-methoxyphenyl)quinolin-4-amine (**17**) showed good activity that tracked with the previously reported trimethoxy derivative.^[17]^ However, the switch from trimethoxy to *meta*-methoxy substitution resulted in a NAK selectivity drop of 6-fold.^[16]^ The switch to the hydroxy moiety (**18**) gave only a slight increase in GAK activity and the same NAK selectivity as the previously reported trimethoxy compound **3**. This trend was repeated in the GAK NanoBRET. The 6-methyl sulfone *meta*-methoxy (**19**) was a weaker GAK binder consistent with previous structure activity relationships observed on this scaffold.^[17]^ The trend observed between **13** and **14** provided the strongest evidence yet that the potential water was being displaced (**Table 2**). A further switch to the nitro substitution with the 6-bromoquinoline **21**, was 10-fold more active than the corresponding 6-trifluoromethyl compound **6** (**Table 3**). This further supports the rational that the 6-trifluoro quinoline does not allow the optimal orientation when compared to the 6-bromoquinoline.

**Table 3.**
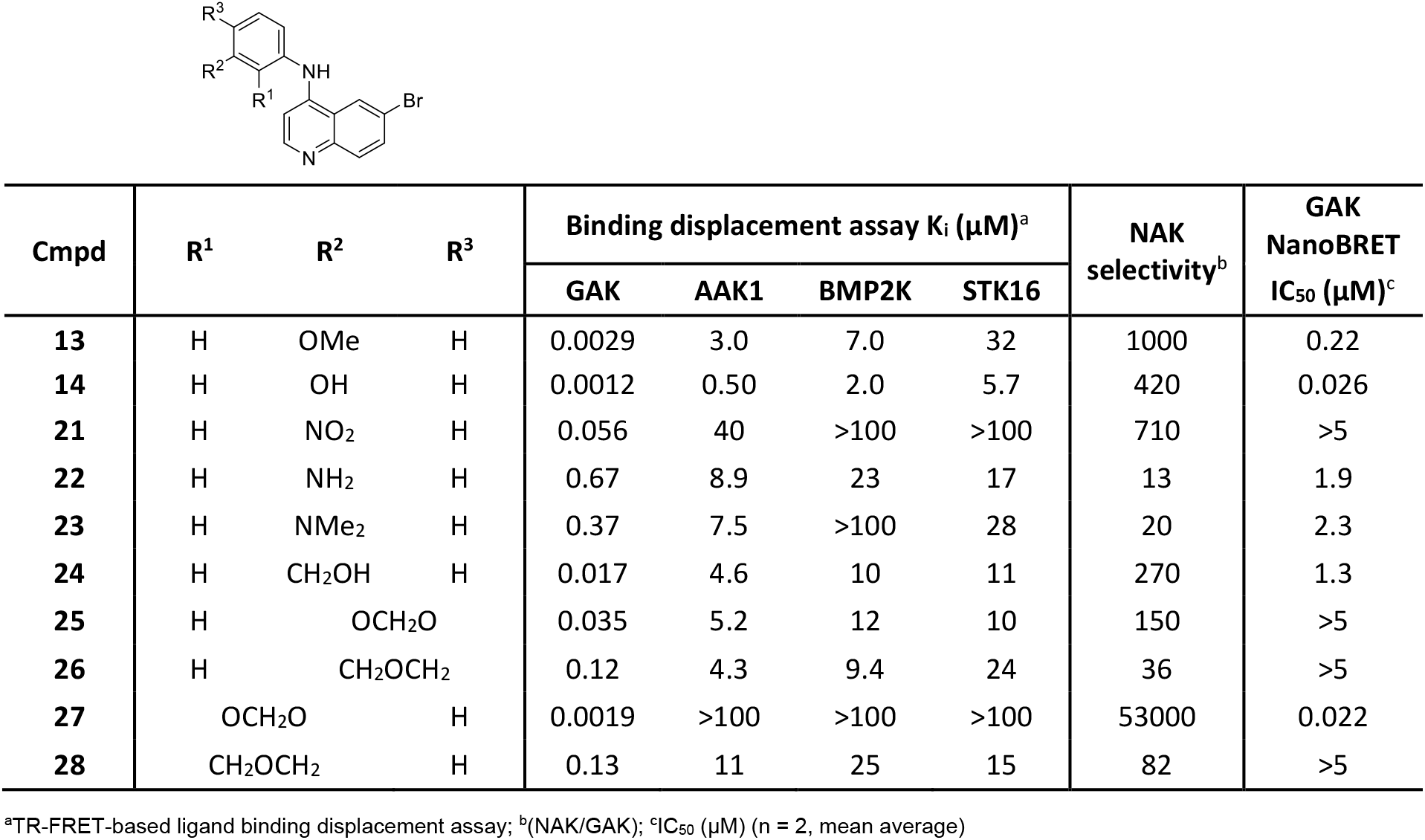
Investigation of water pocket with isosteric replacement of *meta*-methoxy on the 6-bromoquinoline.

However, surprisingly the *meta*-amino group of the 6-bromo compound **22** was 3-fold less potent on GAK compared to **7**. The dimethyl amino aniline derivative **22** was broadly in line with expectations with similar potency to the 6-trifluoromethylquinolines **7** and **8**. The switch from an oxygen atom to a nitrogen atom reduces GAK potency which would suggest that the electronegativity of the oxygen atom is required for the optimal replacement of the water.^[38]^ The *meta*-methanol derivative **24** showed only a limited improvement towards GAK binding compared to the 6-trifluoromethyl (**11**).

We then explored how torsional strain on the methoxy orientation and constrained ring systems would affect the GAK activity profile. The fused ring systems (**25**-**28**) demonstrate how sensitive the water network is and how an island of activity can be achieved. Compounds **25** and **27** demonstrate the same effect as **1** and **2**. The 3,4 connectivity (**25**) vs 2,3 connectivity (**27**) showed a preference for the orthogonal plane angle and an ability to accommodate this in the ATP pocket. However, with **25**, the rotational energy penalty combined with constraints at the 4-position accounts for a 14-fold drop in potency on GAK (**25** vs **27**).

The drop is more severe with the mono-substituted central oxygen which produces two equipotent compounds (**26** & **28**) regardless of positioning (**Figure S1**). The result was the identification of a potent cell active GAK inhibitor (**27**).

We then looked at the conformation of the compounds and the influence this has on the potential water displacement (**Table 4**). Switching from the quinoline *meta*-methoxy aniline (**13**) to quinazoline (**29**) results in a 4-fold penalty, likely related to the flattening of the molecules conformation. The corresponding *meta*-hydroxy analog **30** was equipotent with **29** in GAK binding, likely due to the inability to effectively form the hydrogen bond to replace the water molecule due to the planer character of the compound. Switching back to the 6-bromoquinoline, *meta*-tetrazole **31** with the potential to form multiple hydrogen bonds, mimicked the nitro substitution (**21**) in potency and selectivity despite occupying a large space. The three compounds pentafluorosulfanyl (**32**),^[39]^ *tert*-butoxy (**33**) and the *tert*-butyl (**34**) were likely too large and lipophilic with each demonstrating GAK binding (K_i_) between 2 and 3 micromolar. This highlights the requirement for small, precisely oriented hydrogen bond donors with correct torsional strain to effectively displace the water and form a strong interaction.

**Table 4.**
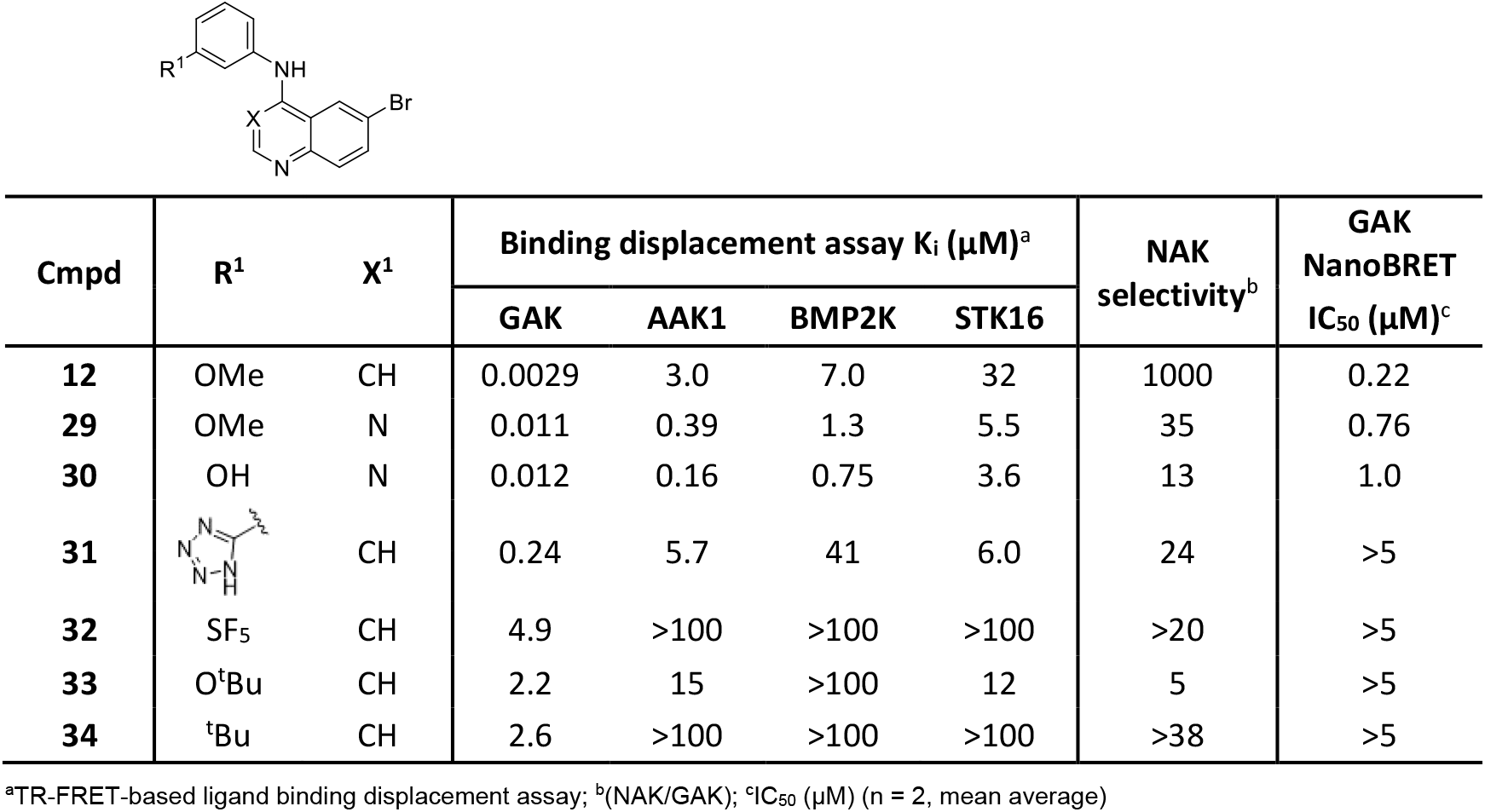
Investigation of water pocket with isoteric replacement of *meta*-methoxy group and different hinge binders.

In order to rationalize the results observed (**Table 1**-**4**) we modelled key compounds SGC-GAK-1, **13**, **14**, and **25**-**28** in the GAK ATP competitive active site with Glide module of Schrödinger Maestro suite (**Figure S1**).^[40]^ We observed through docking that unlike SGC-GAK-1, the *meta*-methoxy derivative (**13**) was not able to form an interaction with the catalytic lysine (Lys69). This runs counter to the observed GAK binding where SGC-GAK-1 and **13** have near equipotency despite this interaction deficit, suggesting other factors including the water network.

The docking of quinazoline **29** demonstrated a lack of the correct orientation to form optimal binding to GAK (**Figure S1**). The docked *meta*-hydroxy (**14**) proposed direct interaction with the alcohol altering the through molecule orientation to form a rarely reported sigma hole interaction (halogen bond) directly with the 6-position bromine mediated through a water molecule bound next to the carbonyl from Leu46 (**Figure S2**).^[41]^ During our investigations this result was not observed with any other analog. The 5-membered dioxane derivative **27** demonstrated the ability to form an optimal fit while still displacing the water (**Figure S1**). Surprisingly none of the other closely related derivatives **25**, **26** and **28** were able to achieve this (**Table 3**).

The WaterMap simulations demonstrate the ability of the most potent GAK inhibitors in the NanoBRET SGC-GAK-1, **13**, **30** and **27** to displace the high energy water present in the hydrophobic pocket. The displacement of this high energy water provides a significant boost of binding affinity. SGC-GAK-1 was able to displace the water and interact with the catalytic lysine (**Figure 6A**). The *meta*-methoxy derivative **13** was able to interact with the catalytic lysine but preferentially chose to displace the water (15/15 simulations) (**Figure 6B**). The WaterMap simulation also demonstrated that the quinazoline **30** had a more flexible conformation and hence was able to only partially displace the water molecule (**Figure 6C**). The 5-member dioxane number was set up in the correct orientation to displace the water molecule (**Figure 6D**). Interestingly, while **25**, **26** and **28** were good binders they were not well set up to displace the high energy water or form key interactions in the ATP binding site of GAK (**Figure S1**).

**Figure 6.**
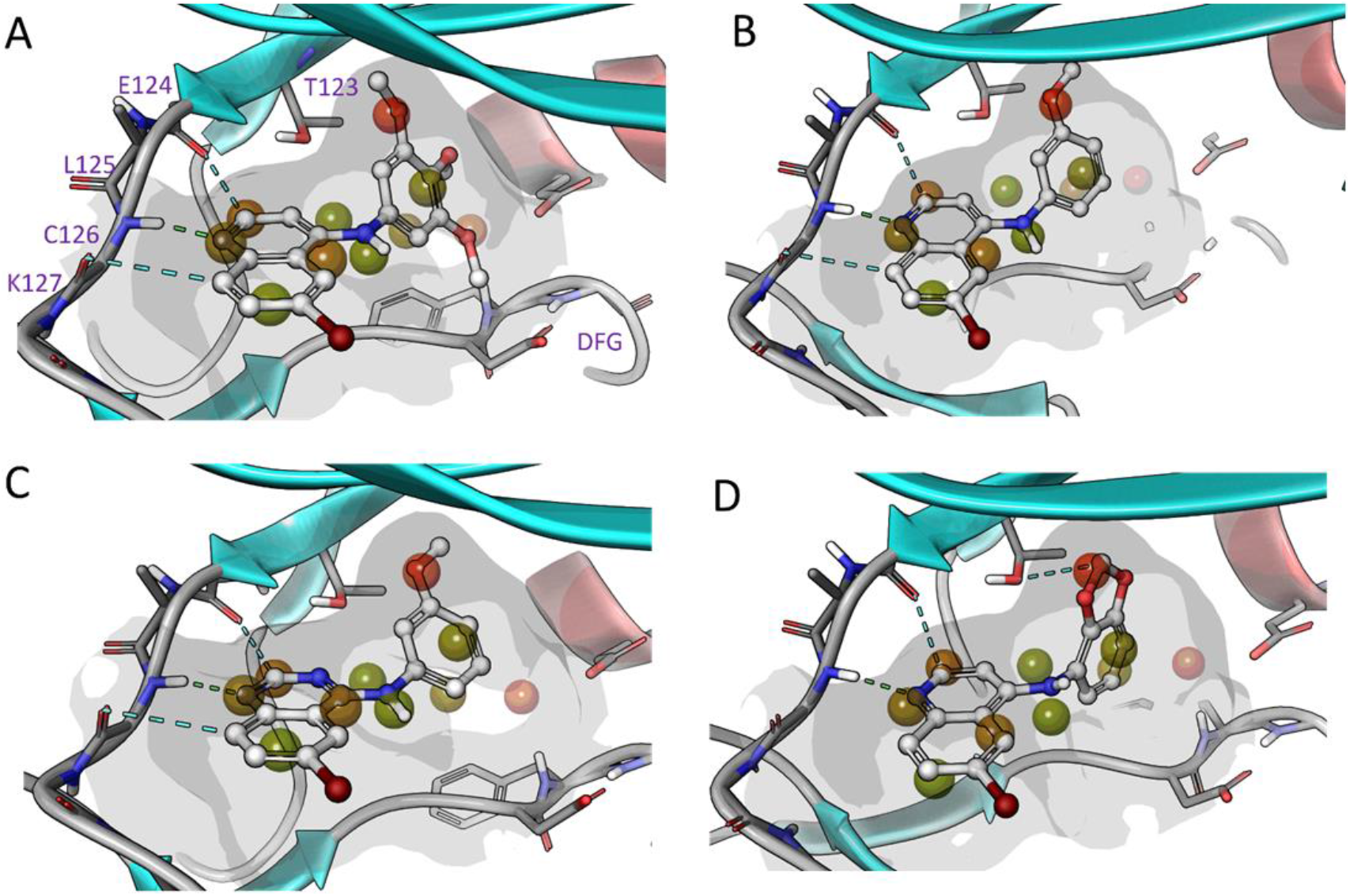
Main WaterMap hydration sites overlaid with selected ligands A - SGC-GAK-1, B - **13**, C - **30** and D - **27** in the GAK ATP binding domain.

In order to investigate the bound water hypothesis further and to further elucidate what constitutes a high and low energy water within the GAK kinase domain we searched for ‘dry areas’ within the ATP binding pocket. These are portions of the receptor active site that are so unfavourable for water molecules that a void is formed there. These are uncommon but have been theoretically and experimentally been observed.^[42]^

The WaterMap analysis of APO structure of GAK (**Figure 7A**) and ligand binding form (**Figure 7B**) of GAK demonstrated that water bound to the high energy pocket had a low relative occupancy of 0.42 in case of APO and 0.53 in case of gefitinib bound, respectively. These low occupancies highlight that there is space, but the positioning of the water is sub-optimal even though a pocket of space available.

**Figure 7.**
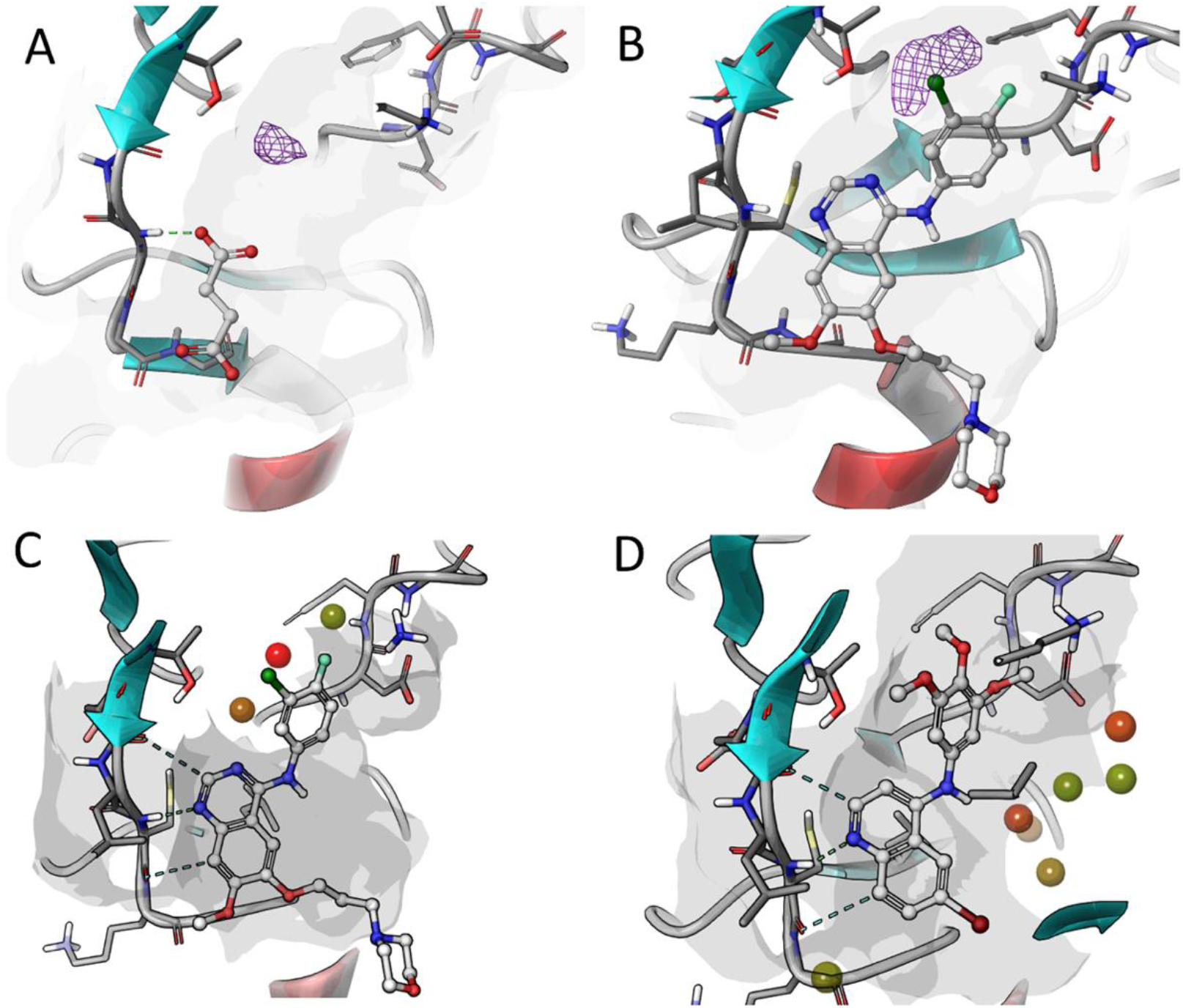
A - GAK APO structure WaterMap (PDB:4C38); B - showing dewetted area extracted from watermap simulation overlaid with gefitinib (PDB:5Y80); C - WaterMap simulation with gefinitib bound to GAK (PDB:5Y80); D - WaterMap simulation with SGC-GAK-1 bound to GAK (PDB:5Y80).

Hydration site analysis with bound gefitinib (**Figure 7C**) showed the pocket present but is only partly occupied by the aniline portion of gefitinib. When gefitinib is switched for SGC-GAK-1 (**Figure 7D**) the hydration site analysis indicates that all the sites are occupied with ligand substituent arms on the trimethoxyaniline portion of the molecule. This provides further explanation for high affinity and enhanced specificity of SGC-GAK-1 and the current series of ligands. An interesting feature of the GAK ATP binding site is that Thr123 is not able to occupy that space at roof of the back pocket which is the origin of this ‘dry’ area.

The observed torsional effects of the aniline to quin(az)oline ring system observed with **13** vs **29** were further explored by solving a series of small molecule crystal structures of **13, 14, 25, 27** and **29** (**Figure 8**). The chloride salts of **13**, **14**, **25** and **27** crystallized and as expected, hydrogen bonding between the amine (and alcohol (**13**)), amine donors and chloride counter-ion are the dominant intermolecular interactions within these structures, leading to simple hydrogen bonded 1D chains in each case. Additionally, **13** and **25** crystallize as hydrates with disordered water molecules in channels parallel to the a- and c-axes respectively resulting in more complex 3D hydrogen bonded networks. Only **29** crystallizes as a pure substance and also forms 1D hydrogen bonded chains *via* the aniline and quinazoline 1-position nitrogen. Structures **13** and **29** crystallize with two independent molecules in the asymmetric unit related by approximate inversion in both cases. The C-N-C-C-C torsion angles of **12**, **13**, **25, 27** and **29** occur in narrow range between ±32.4(8)° - ±61.62(18)° with **12** and **27** displaying torsion angles >±50° The functional groups of the aniline moiety are positioned above the rotation axis of the aniline ring of **12** and **13** whereas they are below the axis of rotation in **25**, **27** and **29**.

**Figure 8.**
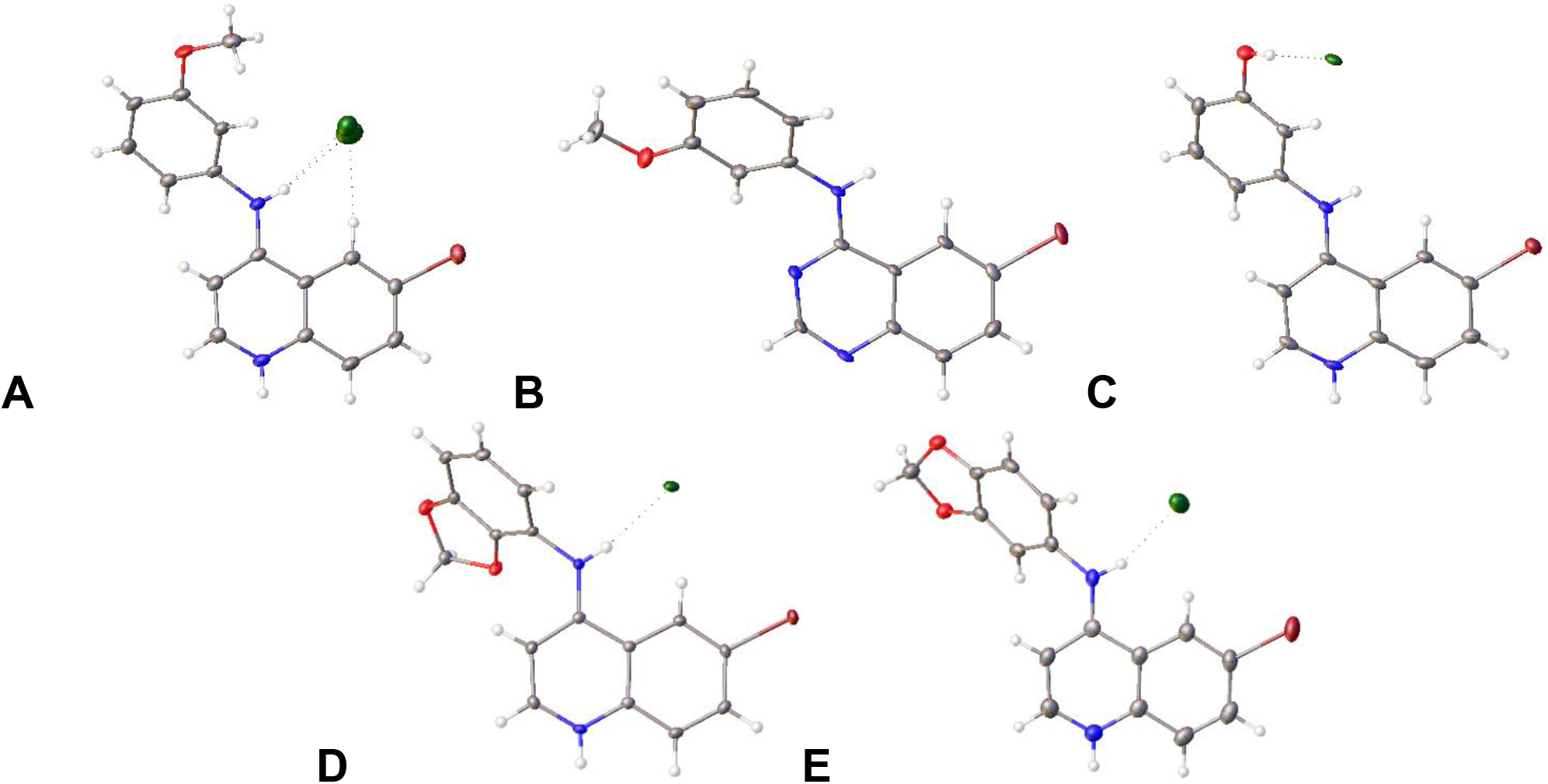
Crystal structures of **13** (A), **29** (B), **14** (C), **27** (D) and **25** (E). ADP ellipsoids are displayed at 50% probability. Counter ions are shown but solvent molecules are not shown for clarity.

## Discussion

The tractability of protein kinases is well known, with more than 50 inhibitors targeting the ATP binding site of kinases approved for use in the clinic.^[43]^ However, most of these drugs leverage the conservation of the ATP binding pocket across kinases to increase their clinical efficacy.^[31]^ These multi-kinase inhibitors while effective would not be well suited outside oncology indications. Development of kinase inhibitors for new treatments outside of oncology will require inhibitors with significantly improved potency and selectivity profiles.^[44]^ Novel approaches to tackle this issue are urgently required in order to effectively develop highly selective and potent kinase inhibitors to target the conserved ATP binding site outside oncology indications.

Several tools to achieve this aim are available with binding assays enabling rapid, accurate and robust method to assess potency and potentially wider selectivity of ATP-competitive kinase inhibitors.^[45–46]^ These ligand binding displacement assays also provide an accepted direct measurement of kinase inhibition in drug optimization of ATP binding site inhibitors.^[45]^ This is a particularly acute point in the case of more neglected kinases such as GAK where there are currently no robust and validated enzyme activity assays.^[47]^

The water network is not a novel concept but has so far attracted limited attention. This is partially due to the fact that targeting and predicting the water network is difficult and, in some cases, not precise enough. WaterMap and other solvent prediction systems have attempted to bridge this gap while other efforts including the *KILFS* database have attempted to map the ATP binding sites across the kinome.^[48]^ Direct utilization of WaterMap has been demonstrated to enable, assess and design selectivity within PI3K subtypes (α, β, γ, and δ) which have highly similar ATP binding sites. The critical role of water molecules in molecular recognition is under recognized and could provide a useful rationalization to drive down potency and improve selectivity.^[49]^

WaterMap analysis of the GAK ATP binding site suggested that a coordinated water network in the protein pocket spans the region proximal the aniline and the 6-position of the ring system. However, one poorly coordinated higher energy water molecule within this network is able to be targeted and displaced. The effect of this displacement leads up to a 10-fold boost due to entropic and enthalpic contributions to the free energy of binding (**Figures 4** - **6**). These models suggest that a water network within the GAK active site plays a critical role in defining the relative affinity of quinoline ligands. Extension of this model to other NAK family members demonstrates why the 4-anilinoquinoline is significantly more potent on GAK compared to the other NAK family members that lack that higher energy water molecule in the lipophilic pocket of the ATP binding site within GAK (**Figure 3** & **4**). The influence of different substituents on the preferred fragment pose was analyzed by various computational approaches. The orthogonal off-targets between **1** and **JMC 2015 - 12g** led us to postulate that the replacement of water molecules results in different flipped binding modes between the two scaffolds (**Figure 9A-B**). This was supported by several computational studies in the literature,^[49–50]^ in addition to a series of oxindole derivatives demonstrating selective DYRK inhibition by a water induced flipped binding mode.^[51]^ Our observations were further bolstered by a recently solved co-crystal structure of gefitinib in GAK showing the same high energy water being displaced (**Figure 9C**).^[52]^

**Figure 9.**
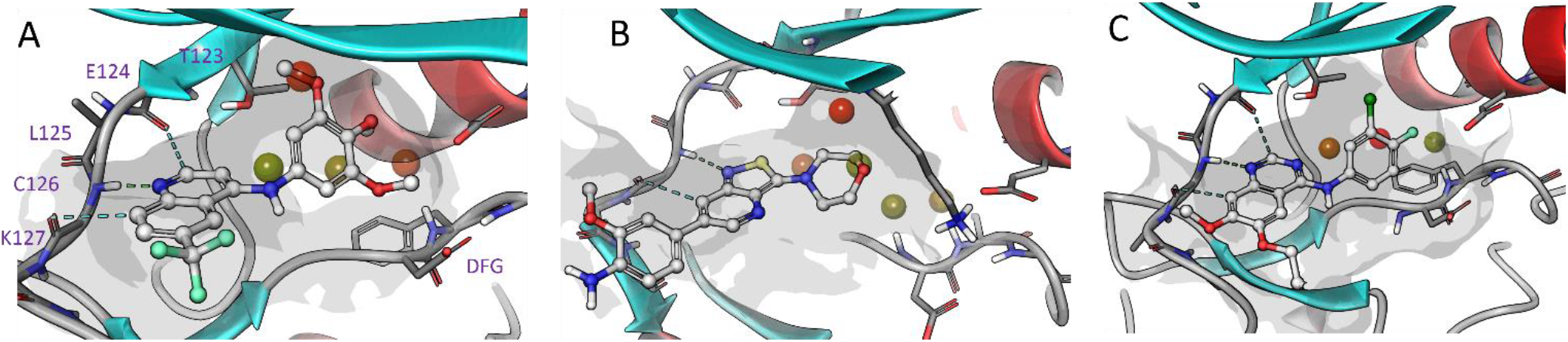
Selected WaterMap hydration sites overlaid with of A: **1** docked (PDB:5Y80), B: **JMC 2015 - 12g** retained at adenosine binding site of GAK (PDB:4Y8D) and C: Gefitinib retained at adenosine binding site (PDB:5Y80).

In summary, we have identified and optimized an island of activity within the ATP binding pocket of GAK by displacement of a high energy water within the lipophilic pocket. The water molecules that occupy the ligand binding pocket prior to small-molecule binding to a protein pocket play a significant role and can be considered as principal source of binding energy. These water molecules occupy certain hydration sites inside the pocket and are either energetically favourable or unfavourable in comparison to the bulk water. If the total energy of a hydration site is positive e.g. it is unfavourable, the replacement of this water will result in a binding affinity boost. A hydration site may be high in energy for multiple reasons, for example if a water is not able to form hydrogen bonds within the site enthalpic penalty results. Entropic penalty can exist when degrees of freedom are unfavourable for water compared to bulk waters. It is also common that a pocket water is transformed to be unfavourable if it is trapped or the degrees of freedom are limited when a small molecule is introduced to the binding site.

In this study we have used WaterMap from Schrödinger to conduct hydration site analysis for family NAK kinases in order to explain ligand affinity further than simple crude scoring functions used in straightforward docking can achieve.^[42,53–56]^ The WaterMap software conducts statistical mechanics based on short molecular dynamics. This simulation describes the thermodynamic properties to estimate the energies of the hydration sites. This estimation is usually computed for the pocket without ligand to evaluated possible hydrations sites but can be also calculated with ligand. However, such short simulations keeping the protein rigid is highly dependent on the conformation of the protein and results are less valid in the case of arbitrary changes in protein conformations. Fortunately, the case of the NAK family kinases is well suited for hydration site analysis thus binding conformations are mapped by several high-resolution x-ray structures resulting in good overall convergence of the ligand dockings.

This method can be used to further optimize GAK inhibitors for potential *in vitro* and *in vivo* usage.^[20,57]^ Extension of this model to other NAK family members or more distantly related kinases could lead to computational models that predict wider kinome selectivity.

## Experimental Section

### Modelling method

Molecular modelling was performed using Schrödinger Maestro software package (Small-Molecule Drug Discovery Suite 2018-4, Schrödinger, LLC, New York, NY, 2018) Prior to docking simulations structures of small molecules were prepared using and the LigPrep module of Schrodinger suite employing OPLS3e force field.^[55]^ In the case of GAK and other NAK family kinases there are a number of co-crystal structures available representing various ligand binding conformations, showing flexibility in the position of so-called the p-loop and C-helix region. Suitable docking templates were searched using LPDB module of Schrödinger package and carrying out visual inspection of available experimental structures with assistance of LiteMol plug-in available at website of UniProt database. Selected coordinates (PDB: 4Y8D and 5Y7Z) have been co-crystallized with at resolution of 2.1Å and 2.5Å respectively with a small molecule inhibitors.^[26,52]^ The PDB structure of GAK was H-bond optimized and minimized using standard protein preparation procedure of Schrödinger suite. The ligand docking was performed using SP settings of Schrodinger docking protocol with softened vdw potential (scaling 0.6). In order to improve convergence of docking poses a hydrogen bond constraint to mainchain NH of hinge residue was required, as experimentally observed in the case of quinoline/quinolizine scaffolds (for example NH of C126 in the case of GAK). The grid box was centered using coordinate center of the core structure of corresponding x-ray ligand as template. Graphical illustrations were generated using, Maestro, and PyMOL software of Schrödinger.

### Hydration Site Analysis

Hydration site analysis calculated with WaterMap (Schrödinger Release 2018-4: WaterMap, Schrödinger, LLC, New York, NY, 2018.). The structure of GAK (PDB: 4Y8D and 5Y7Z) was prepared with Protein Preparation Wizard (as above).^[27,52]^ Water molecules were analyzed within 6 Å from the docked ligand, and the 2 ns simulation was conducted with OPLS3e force field.^[55]^

### Cellular NanoBRET target engagement assay

Screening was performed as previously described.^[17]^

### Ligand binding displacement assays

Screening was performed as previously described,^[17]^

Constructs used: AAK1 - AAKA-p051; BMPK2K - BMP2KA-p031; GAK - GAKA-p059; STK16 - STK16A-p016.^[17]^

### Chemistry

General procedure for the synthesis of 4-anilinoquin(az)olines: 4-chloroquin(az)oline derivative (1.0 eq.) and aniline derivative (1.1 eq.) were suspended in ethanol (10 mL) and refluxed for 18h. The crude mixture was purified by flash chromatography using EtOAc:hexane followed by 1-5 % methanol in EtOAc; After solvent removal under reduced pressure, the product was obtained as a solid or recrystallized from ethanol/water. Compounds **1**-**4** were prepared as previously described.^[17]^

**3-((6-(trifluoromethyl)quinolin-4-yl)amino)phenol (5)** was obtained as a mustard solid (114 mg, 0.376 mmol, 58 %). MP 166-168 °C; ^1^H NMR (400 MHz, DMSO-*d*_6_) δ 11.09 (s, 1H), 10.05 (s,1H), 9.32 (s, 1H), 8.56 (d, *J* = 6.6 Hz, 1H), 8.39 – 8.11 (m, 2H), 7.32 (t, *J* = 8.0 Hz, 1H), 7.13 – 6.61 (m, 4H). ^13^C NMR (100 MHz, DMSO-*d*_6_) δ 158.8, 154.3, 145.5, 142.0, 138.3, 130.6, 128.4 (d, *J* = 3.5 Hz), 126.3 (q, *J* = 32.7 Hz), 125.3, 123.4, 122.5 (d, *J* = 5.2 Hz), 117.0, 115.1, 114.3, 111.8, 101.4. HRMS *m/z* [M+H]^+^ calcd for C_16_H_12_N_2_OF_3_: 305.0902, found 305.0892, LC *t*_R_ = 3.38 min, > 98 % Purity.

***N*-(3-nitrophenyl)-6-(trifluoromethyl)quinolin-4-amine (6)** (179 mg, 0.538 mmol, 83%) ^1^H NMR (400 MHz, DMSO-*d*_6_) δ 11.84 (s, 1H), 9.59 – 9.33 (m, 1H), 8.69 (d, *J* = 7.0 Hz, 1H), 8.38 (dd, *J* = 4.5, 2.4 Hz, 2H), 8.32 (dd, *J* = 9.0, 1.7 Hz, 1H), 8.24 (ddd, *J* = 8.3, 2.3, 1.0 Hz, 1H), 8.02 (ddd, *J* = 8.0, 2.1, 1.0 Hz, 1H), 7.86 (t, *J* = 8.1 Hz, 1H), 7.10 (d, *J* = 7.0 Hz, 1H). ^13^C NMR (100 MHz, DMSO-*d*_6_) δ 155.2, 148.6, 144.6, 140.3, 138.4, 131.4 (d, *J* = 17.2 Hz), 129.4 (d, *J* = 3.4 Hz), 127.8, 127.0 (q, *J* = 32.9 Hz), 125.1, 123.0 (q, *J* = 4.1 Hz), 122.4, 122.0 (d, *J* = 26.5 Hz), 120.0, 117.1, 101.6. HRMS *m/z* [M+H]^+^ calcd for C_16_H_11_N_3_O_2_F_3_: 334.0803, found 334.0793, LC *t*_R_ = 3.78 min, > 98 % Purity.

***N*^1^-(6-(trifluoromethyl)quinolin-4-yl)benzene-1,3-diamine (7) 6** (150 mg) was treated with palladium on carbon under hydrogen for 18 h and purified by flash chromatography using EtOAc:hexane followed by 2 % methanol in EtOAc. The solvent was removed under reduced pressure, the product was obtained as a dark yellow solid (94 mg, 0.311 mmol, 69%). ^1^H NMR (400 MHz, DMSO-*d*_6_) δ 10.79 (s, 1H), 8.86 – 8.72 (m, 1H), 8.47 (d, *J* = 6.9 Hz, 1H), 8.09 (dd, *J* = 8.5, 1.2 Hz, 1H), 7.98 (ddd, *J* = 8.4, 6.9, 1.2 Hz, 1H), 7.75 (ddd, *J* = 8.4, 6.9, 1.2 Hz, 1H), 7.18 (t, *J* = 8.0 Hz, 1H), 6.78 (d, *J* = 6.9 Hz, 1H), 6.71 – 6.38 (m, 3H), 5.48 (s, 2H). ^13^C NMR (100 MHz, DMSO-*d*_6_) δ 154.8, 150.4, 142.6, 138.7, 137.8, 133.6, 130.2, 126.7, 123.6, 120.5, 117.0, 113.0, 112.1, 110.2, 99.9. HRMS *m/z* [M+H]^+^ calcd for C_16_H_13_N_3_F_3_: 304.1062, found 304.1051, LC *t*_R_ = 3.20 min, > 98 % Purity.

**^*1*^*N*,^*1*^*N*-dimethyl-3-*N*-[6-(trifluoromethyl)quinolin-4-yl]benzene-1,3-diamine (8)** was obtained as a dark yellow solid (163 mg, 0.492 mmol, 76%). MP decomp >160 °C; ^1^H NMR (400 MHz, DMSO-*d*_6_) δ 11.37 (s, 1H), 9.36 (dt, *J* = 1.9, 1.0 Hz, 1H), 8.55 (d, *J* = 7.1 Hz, 1H), 8.46 – 8.08 (m, 2H), 7.45 – 7.25 (m, 1H), 6.87 (d, *J* = 7.1 Hz, 1H), 6.86 – 6.75 (m, 2H), 6.76 – 6.60 (m, 1H), 2.94 (s, 6H). ^13^C NMR (101 MHz, DMSO-*d*_6_) δ 155.6, 151.6, 143.8, 140.3, 137.5, 130.3, 129.2 (d, *J* = 3.4 Hz), 126.6 (q, *J* = 32.7 Hz), 125.1, 122.6 (q, *J* = 4.1 Hz), 121.9, 116.6, 112.3, 111.6, 108.7, 101.2, 40.0 (s, 2C). HRMS *m/z* [M+H]^+^ calcd for C_18_H_17_N_3_F_3_: 332.1375, found 332.1366, LC *t*_R_ = 4.02 min, > 98 % Purity.

***N*-(3-nitrophenyl)quinolin-4-amine (9)** was obtained as a yellow solid (217 mg, 0.816 mmol, 89%). MP >250 °C; ^1^H NMR (400 MHz, DMSO-*d*_6_) δ 11.39 (s, 1H), 8.94 (dd, *J* = 8.6, 1.2 Hz, 1H), 8.61 (d, *J* = 6.9 Hz, 1H), 8.38 (t, *J* = 2.2 Hz, 1H), 8.22 (ddd, *J* = 8.3, 2.3, 1.0 Hz, 1H), 8.17 (dd, *J* = 8.6, 1.2 Hz, 1H), 8.08 – 8.00 (m, 2H), 7.87 – 7.81 (m, 2H), 7.04 (d, *J* = 6.8 Hz, 1H). ^13^C NMR (101 MHz, DMSO-*d*_6_) δ 154.6, 148.6, 143.2, 138.9, 138.4, 134.0, 131.4, 131.2, 127.3, 124.0, 121.5, 120.4, 119.8, 117.6, 100.4. HRMS *m/z* [M+H]^+^ calcd for C_15_H_12_N_3_O_2_: 266.0930, found 266.0920, LC *t*_R_ = 2.97 min, > 98 % Purity.

***N*^*1*^,*N*^*1*^-dimethyl-*N*^*3*^-(quinolin-4-yl)benzene-1,3-diamine (10)** was obtained as a dark yellow solid (188 mg, 0.715 mmol, 78%) MP >250 °C; ^1^H NMR (400 MHz, DMSO-*d*_6_) δ 10.98 (s, 1H), 8.85 (dd, *J* = 8.7, 1.2 Hz, 1H), 8.47 (d, *J* = 7.0 Hz, 1H), 8.11 (dd, *J* = 8.6, 1.2 Hz, 1H), 8.00 (ddd, *J* = 8.4, 6.9, 1.2 Hz, 1H), 7.77 (ddd, *J* = 8.3, 6.9, 1.2 Hz, 1H), 7.35 (td, *J* = 7.8, 0.9 Hz, 1H), 6.80 (d, *J* = 7.0 Hz, 1H), 6.78 – 6.44 (m, 3H), 2.94 (s, 6H). ^13^C NMR (101 MHz, DMSO-*d*_6_) δ 155.1, 151.6, 142.4, 138.3, 137.9, 133.7, 130.2, 126.8, 123.7, 120.2, 117.0, 112.5, 111.3, 109.0, 100.0, 40.0 (s, 2C). HRMS *m/z* [M+H]^+^ calcd for C_17_H_18_N_3_: 264.1501, found 264.1493, LC *t*_R_ = 3.09 min, > 98 % Purity.

**(3-{[6-(trifluoromethyl)quinolin-4-yl]amino}phenyl)methanol (11)** was obtained as a light yellow solid (151 mg, 0.473 mmol, 73%). MP 152-154 °C; ^1^H NMR (400 MHz, DMSO-*d*_6_) δ 10.94 (s, 1H), 9.28 (d, *J* = 1.9 Hz, 1H), 8.57 (d, *J* = 6.6 Hz, 1H), 8.26 (d, *J* = 8.9 Hz, 1H), 8.19 (dd, *J* = 8.9, 1.8 Hz, 1H), 7.49 (t, *J* = 7.7 Hz, 1H), 7.43 (t, *J* = 1.8 Hz, 1H), 7.32 (ddt, *J* = 10.4, 7.6, 1.2 Hz, 2H), 6.90 (d, *J* = 6.6 Hz, 1H), 5.42 (s, 1H), 4.57 (s, 2H).^13^C NMR (100 MHz, DMSO-*d*_6_) δ 153.6, 146.5, 144.7, 143.0, 137.6, 129.5, 128.0 (d, *J* = 2.7 Hz), 126.2 (q, *J* = 32.7 Hz), 125.4, 124.7, 124.3, 122.4, 122.3 (d, *J* = 4.3 Hz), 122.3, 117.3, 101.3, 62.4. HRMS *m/z* [M+H]^+^ calcd for C_17_H_14_N_2_OF_3_: 319.1058, found 319.1048, LC *t*_R_ = 3.26 min, > 98 % Purity.

**(2-{[6-(trifluoromethyl)quinolin-4-yl]amino}phenyl)methanol (12)** was obtained as a grey/yellow solid (132 mg, 0.416 mmol, 64%). MP 120-122 °C; ^1^H NMR (400 MHz, DMSO-*d*_6_) δ 10.18 (s, 1H), 9.18 (s, 1H), 8.48 (d, *J* = 6.1 Hz, 1H), 8.33 – 7.85 (m, 2H), 7.84 – 7.56 (m, 1H), 7.56 – 6.93 (m, 3H), 6.27 (d, *J* = 6.1 Hz, 1H), 5.33 (s, 1H), 4.49 (s, 2H). ^13^C NMR (100 MHz, DMSO-*d*_6_) δ 152.9, 148.9, 145.7, 139.2, 135.3, 128.4, 128.2, 127.5, 127.0, 126.9 – 126.4 (m), 125.6 (d, *J* = 4.9 Hz), 125.3, 122.9, 121.8 (q, *J* = 4.3 Hz), 117.4, 101.3, 59.0. HRMS *m/z* [M+H]^+^ calcd for C_17_H_14_N_2_OF_3_: 319.1058, found 319.1048, LC *t*_R_ = 3.24 min, > 98 % Purity.

**6-bromo-*N*-(3-methoxyphenyl)quinolin-4-amine (13)** was obtained as a a beige solid (151 mg, 0.458 mmol, 74 %). MP 188-190 °C; ^1^H NMR (400 MHz, DMSO-*d*_6_) δ 11.07 (s, 1H), 9.16 (d, *J* = 2.0 Hz, 1H), 8.52 (d, *J* = 7.0 Hz, 1H), 8.17 (dd, *J* = 9.0, 2.0 Hz, 1H), 8.08 (d, *J* = 9.0 Hz, 1H), 7.48 (t, *J* = 8.4 Hz, 1H), 7.31 – 7.03 (m, 2H), 7.00 (ddd, *J* = 8.3, 2.4, 1.0 Hz, 1H), 6.89 (d, *J* = 6.9 Hz, 1H), 3.81 (s, 3H). ^13^C NMR (100 MHz, DMSO-*d*_6_) δ ^13^C NMR (101 MHz, DMSO-*d*_6_) δ 160.3, 153.9, 143.0, 138.2, 137.4, 136.6, 130.8, 126.2, 122.5, 119.8, 118.6, 117.1, 113.1, 110.9, 100.7, 55.4. HRMS *m/z* [M+H]^+^ calcd for C_16_H_14_N_2_BrO: 329.0286, found 329.0289, LC *t*_R_ = 3.63 min, > 98 % Purity.

**3-((6-bromoquinolin-4-yl)amino)phenol (14)** was obtained as a yellow solid (138 mg, 0.437 mmol, 71 %). MP >300 °C; ^1^H NMR (400 MHz, DMSO-*d*_6_) δ 11.00 (s, 1H), 10.03 (s, 1H), 9.14 (d, *J* = 2.0 Hz, 1H), 8.51 (d, *J* = 7.0 Hz, 1H), 8.15 (dd, *J* = 9.0, 2.0 Hz, 1H), 8.06 (d, *J* = 9.0 Hz, 1H), 7.42 – 7.29 (m, 1H), 7.10 – 6.66 (m, 4H). ^13^C NMR (100 MHz, DMSO-*d*_6_) δ 159.2, 154.4, 143.3, 138.3, 137.7, 137.0, 131.1, 126.6, 122.8, 120.2, 119.0, 115.9, 115.2, 112.5, 101.0. HRMS *m/z* [M+H]^+^ calcd for C_15_H_11_BrN_2_O: 315.0132, found 315.0124, LC *t*_R_ = 3.01 min, > 98 % Purity.

***N*-(3-methoxyphenyl)quinolin-4-amine (15)** was obtained as a grey solid (191 mg, 0.761 mmol, 81 %). MP 198-200 °C; ^1^H NMR (400 MHz, DMSO-*d*_6_) δ 11.08 (s, 1H), 8.88 (dd, *J* = 8.6, 1.2 Hz, 1H), 8.50 (d, *J* = 7.0 Hz, 1H), 8.13 (dd, *J* = 8.6, 1.2 Hz, 1H), 8.02 (ddd, *J* = 8.4, 6.9, 1.2 Hz, 1H), 7.79 (ddd, *J* = 8.4, 6.9, 1.2 Hz, 1H), 7.54 – 7.40 (m, 1H), 7.31 – 7.04 (m, 2H), 7.04 – 6.89 (m, 1H), 6.85 (d, *J* = 6.9 Hz, 1H), 3.81 (s, 3H). ^13^C NMR (100 MHz, DMSO-*d*_6_) δ 160.3, 154.9, 142.6, 138.4, 138.2, 133.8, 130.7, 127.0, 123.8, 120.2, 117.4, 117.1, 113.1, 111.2, 100.0, 55.4. HRMS *m/z* [M+H]^+^ calcd for C_16_H_15_N_2_O: 251.1184, found 251.1175, LC *t*_R_ = 3.17 min, > 98 % Purity.

**3-(quinolin-4-ylamino)phenol (16)** was obtained as a green/yellow solid (74 mg, 0.312 mmol, 34 %). MP decomp. >230 °C; ^1^H NMR (400 MHz, DMSO-*d*_6_) δ 10.97 (s, 1H), 10.03 (s, 1H), 8.84 (dd, *J* = 8.7, 1.1 Hz, 1H), 8.50 (d, *J* = 7.0 Hz, 1H), 8.11 (dd, *J* = 8.5, 1.2 Hz, 1H), 8.01 (ddd, *J* = 8.4, 6.9, 1.2 Hz, 1H), 7.77 (ddd, *J* = 8.3, 6.9, 1.2 Hz, 1H), 7.48 – 7.19 (m, 1H), 6.91 – 6.80 (m, 3H). ^13^C NMR (100 MHz, DMSO-*d*_6_) δ 158.8, 155.0, 142.5, 138.2, 138.0, 133.8, 130.6, 126.9, 123.8, 120.1, 117.1, 115.7, 114.7, 112.3, 99.9. HRMS *m/z* [M+H]^+^ calcd for C_15_H_13_N_2_O: 237.1028, found 237.1019, LC *t*_R_ = 2.66 min, > 98 % Purity.

**6-methoxy-*N*-(3-methoxyphenyl)quinolin-4-amine (17)** was obtained as a beige/yellow solid (148 mg, 0.527 mmol, 68 %). MP 170-172 °C; ^1^H NMR (400 MHz, DMSO-*d*_6_) δ 10.93 (s, 1H), 8.41 (d, *J* = 6.9 Hz, 1H), 8.28 (d, *J* = 2.6 Hz, 1H), 8.06 (d, *J* = 9.3 Hz, 1H), 7.66 (dd, *J* = 9.2, 2.6 Hz, 1H), 7.52 – 7.40 (m, 1H), 7.28 – 7.03 (m, 2H), 6.99 (ddd, *J* = 8.4, 2.4, 1.1 Hz, 1H), 6.84 (d, *J* = 6.8 Hz, 1H), 4.00 (s, 3H), 3.81 (s, 3H). ^13^C NMR (100 MHz, DMSO-*d*_6_) δ 160.3, 158.0, 153.8, 140.5, 138.6, 133.5, 130.7, 125.4, 121.9, 118.5, 117.4, 112.8, 111.1, 103.0, 99.8, 56.6, 55.4. HRMS *m/z* [M+H]^+^ calcd for C_17_H_17_N_2_O_2_: 281.1290, found 281.1281, LC *t*_R_ = 3.58 min, > 98 % Purity.

**3-((6-methoxyquinolin-4-yl)amino)phenol (18)** was obtained as a beige solid (78 mg, 0.294 mmol, 38 %). MP >300 °C; ^1^H NMR (400 MHz, DMSO-*d*_6_) δ 10.79 (s, 1H), 9.99 (s, 1H), 8.40 (d, *J* = 6.9 Hz, 1H), 8.23 (d, *J* = 2.6 Hz, 1H), 8.04 (d, *J* = 9.3 Hz, 1H), 7.65 (dd, *J* = 9.2, 2.5 Hz, 1H), 7.39 – 7.29 (m, 1H), 7.00 – 6.71 (m, 4H), 3.99 (s, 3H). ^13^C NMR (100 MHz, DMSO-*d*_6_) δ 158.7, 157.9, 153.8, 140.5, 138.3, 133.5, 130.6, 125.3, 121.9, 118.4, 115.7, 114.4, 112.3, 102.9, 99.6, 56.5. HRMS *m/z* [M+H]^+^ calcd for C_16_H_15_N_2_O_2_: 267.1134, found 267.1124, LC *t*_R_ = 3.05 min, > 98 % Purity.

**6-methanesulfonyl-*N*-(3-methoxyphenyl)quinolin-4-amine (19)** was obtained as a yellow solid (171 mg, 0.521 mmol, 84 %). MP 271-273 °C; ^1^H NMR (400 MHz, DMSO-*d*_6_) δ 11.61 (s, 1H), 9.55 (d, *J* = 1.8 Hz, 1H), 8.59 (d, *J* = 7.0 Hz, 1H), 8.44 (dd, *J* = 8.9, 1.8 Hz, 1H), 8.31 (d, *J* = 8.9 Hz, 1H), 7.54 – 7.45 (m, 1H), 7.14 – 7.05 (m, 2H), 7.03 (ddd, *J* = 8.4, 2.3, 1.1 Hz, 1H), 6.95 (d, *J* = 7.0 Hz, 1H), 3.81 (s, 3H), 3.44 (s, 3H). ^13^C NMR (100 MHz, DMSO-*d*_6_) δ 160.4, 155.6, 144.4, 140.7, 138.7, 138.0, 130.8, 130.4, 124.9, 121.8, 117.1, 116.8, 113.3, 110.9, 101.5, 55.4, 43.5. HRMS *m/z* [M+H]^+^ calcd for C_17_H_17_N_2_O_3_S: 329.0960, found 329.0953, LC *t*_R_ = 3.08 min, > 98 % Purity.

**3-((6-(methylsulfonyl)quinolin-4-yl)amino)phenol (20)** was obtained as a yellow solid (98 mg, 0.312 mmol, 50 %). MP >300 °C; ^1^H NMR (400 MHz, DMSO-*d*_6_) δ 11.51 (s, 1H), 10.03 (s, 1H), 9.51 (d, *J* = 1.8 Hz, 1H), 8.58 (d, *J* = 7.1 Hz, 1H), 8.43 (dd, *J* = 8.9, 1.8 Hz, 1H), 8.28 (d, *J* = 8.9 Hz, 1H), 7.37 (t, *J* = 8.3 Hz, 1H), 7.09 – 6.66 (m, 4H), 3.43 (s, 3H). ^13^C NMR (100 MHz, DMSO-*d*_6_) δ 158.8, 155.6, 144.2, 140.7, 138.6, 137.8, 130.7, 130.4, 124.9, 121.8, 116.8, 115.4, 114.9, 112.0, 101.3, 43.5. HRMS *m/z* [M+H]^+^ calcd for C_16_H_15_N_2_O_3_S: 315.0803, found 315.0796, LC *t*_R_ = 2.29 min, > 98 % Purity.

**6-bromo-*N*-(3-nitrophenyl)quinolin-4-amine (21)** was obtained as a beige solid (162 mg, 0.470 mmol, 76 %). MP 290-292 °C; ^1^H NMR (400 MHz, DMSO-*d*_6_) δ 11.39 (s, 1H), 9.23 (d, *J* = 2.0 Hz, 1H), 8.62 (d, *J* = 6.9 Hz, 1H), 8.36 (t, *J* = 2.2 Hz, 1H), 8.30 – 8.15 (m, 2H), 8.13 (d, *J* = 9.0 Hz, 1H), 8.00 (ddd, *J* = 8.0, 2.1, 1.0 Hz, 1H), 7.84 (t, *J* = 8.1 Hz, 1H), 7.08 (d, *J* = 6.9 Hz, 1H). ^13^C NMR (100 MHz, DMSO-*d*_6_) δ 153.6, 148.6, 143.5, 138.6, 137.5, 136.7, 131.3, 131.2, 126.4, 122.6, 121.6, 120.2, 119.7, 119.0, 101.1. HRMS *m/z* [M+H]^+^ calcd for C_15_H_11_N_3_O_2_Br: 344.0035, found 344.0032, LC *t*_R_ = 3.48 min, > 98 % Purity.

**1-*N*-(6-bromoquinolin-4-yl)benzene-1,3-diamine (22) 21** (150 mg) was treated with palladium on carbon under hydrogen for 18 h and purified by flash chromatography using EtOAc:hexane followed by 2 % methanol in EtOAc. The solvent was removed under reduced pressure, the product was was obtained as a brown solid (129 mg, 0.410 mmol, 94 %). MP 280-282 °C; ^1^H NMR (400 MHz, DMSO-*d*_6_) δ 11.06 (s, 1H), 8.85 (d, *J* = 8.6 Hz, 1H), 8.58 (d, *J* = 6.9 Hz, 1H), 8.16 – 8.00 (m, 2H), 7.81 (t, *J* = 7.8 Hz, 1H), 7.57 (t, *J* = 8.2 Hz, 1H), 7.34 (d, *J* = 6.0 Hz, 2H), 7.24 (d, *J* = 7.7 Hz, 1H), 6.89 (d, *J* = 6.9 Hz, 1H). ^13^C NMR (100 MHz, DMSO-*d*_6_) δ 154.8, 142.8, 138.2, 138.1, 134.0, 130.9, 127.2, 123.8, 123.6, 121.6, 120.2, 119.8, 117.6, 117.2, 100.0. HRMS *m/z* [M+H]^+^ calcd for C_15_H_13_BrN_3_: 314.0293, found 314.0293, LC *t*_R_ = 2.22 min, > 98 % Purity.

**3-*N*-(6-bromoquinolin-4-yl)-1-*N*,1-*N*-dimethylbenzene-1,3-diamine (23)** was obtained as a yellow/mustard solid (144 mg, 0.421 mmol, 68 %). MP Decomp. >280 °C; ^1^H NMR (400 MHz, DMSO-*d*_6_) δ 10.99 (s, 1H), 9.14 (d, *J* = 2.0 Hz, 1H), 8.48 (d, *J* = 6.9 Hz, 1H), 8.15 (dd, *J* = 9.0, 2.0 Hz, 1H), 8.07 (d, *J* = 9.0 Hz, 1H), 7.39 – 7.27 (m, 1H), 6.84 (d, *J* = 7.0 Hz, 1H), 6.81 – 6.51 (m, 3H), 2.94 (s, 6H). ^13^C NMR (100 MHz, DMSO-*d*_6_) δ 154.2, 151.6, 142.8, 137.7, 137.3, 136.5, 130.2, 126.1, 122.4, 119.7, 118.5, 112.2, 111.38, 108.7, 100.6, 39.98 (2C, s). HRMS *m/z* [M+H]^+^ calcd for C_17_H_17_N_3_Br: 342.0606, found 342.0597, LC *t*_R_ = 3.74 min, > 98 % Purity.

**{3-[(6-bromoquinolin-4-yl)amino]phenyl}methanol (24)** was obtained as a yellow/mustard solid (155 mg, 0.470 mmol, 76 %). MP 132-134 °C; ^1^H NMR (400 MHz, DMSO-*d*_6_) δ 10.93 (s, 1H), 9.13 (d, *J* = 2.0 Hz, 1H), 8.52 (d, *J* = 6.8 Hz, 1H), 8.14 (dd, *J* = 9.0, 1.9 Hz, 1H), 8.06 (d, *J* = 9.0 Hz, 1H), 7.51 (t, *J* = 7.7 Hz, 1H), 7.45 – 7.11 (m, 3H), 6.85 (d, *J* = 6.8 Hz, 1H), 5.41 (s, 1H), 4.58 (s, 2H). ^13^C NMR (100 MHz, DMSO-*d*_6_) δ 153.5, 144.8, 143.6, 138.1, 137.1, 136.2, 129.6, 126.1, 125.1, 123.1, 122.7, 119.7, 118.8, 100.5, 62.3. HRMS *m/z* [M+H]^+^ calcd for C_16_H_14_N_2_OBr: 329.0289, found 329.0283, LC *t*_R_ = 4.20 min, > 98 % Purity.

***N*-(2*H*-1,3-benzodioxol-5-yl)-6-bromoquinolin-4-amine (25)** was obtained as a green solid (144 mg, 0.421 mmol, 68 %). MP 240-242 °C; ^1^H NMR (400 MHz, DMSO-*d*_6_) δ 11.00 (s, 1H), 9.14 (d, *J* = 2.0 Hz, 1H), 8.49 (d, *J* = 7.0 Hz, 1H), 8.15 (dd, *J* = 9.0, 2.0 Hz, 1H), 8.06 (d, *J* = 9.0 Hz, 1H), 7.44 – 7.00 (m, 2H), 6.93 (dd, *J* = 8.2, 2.1 Hz, 1H), 6.73 (d, *J* = 6.9 Hz, 1H), 6.13 (s, 2H). ^13^C NMR (100 MHz, DMSO-*d*_6_) δ 154.5, 148.2, 146.6, 142.8, 137.3, 136.5, 130.6, 126.2, 122.4, 119.7, 119.2, 118.4, 108.9, 106.9, 101.9, 100.4. HRMS *m/z* [M+H]^+^ calcd for C_16_H_12_N_2_O_2_Br: 343.0082, found 343.0079, LC *t*_R_ = 3.49 min, > 98 % Purity.

**6-bromo-*N*-(1,3-dihydro-2-benzofuran-5-yl)quinolin-4-amine (26)** was obtained as a yellow solid (148 mg, 0.433 mmol, 70 %). MP 140-142 °C; ^1^H NMR (400 MHz, DMSO-*d*_6_) δ 11.22 (s, 1H), 9.22 (d, *J* = 2.0 Hz, 1H), 8.49 (d, *J* = 7.0 Hz, 1H), 8.14 (dd, *J* = 9.0, 1.9 Hz, 1H), 8.08 (d, *J* = 9.0 Hz, 1H), 7.48 (d, *J* = 7.8 Hz, 1H), 7.42 (d, *J* = 2.0 Hz, 1H), 7.37 (dd, *J* = 8.0, 1.9 Hz, 1H), 6.80 (d, *J* = 7.0 Hz, 1H), 5.05 (s, 4H). ^13^C NMR (100 MHz, DMSO-*d*_6_) δ 154.6, 143.3, 141.5, 138.8, 137.7, 137.0, 136.6, 126.7, 125.0, 123.0, 122.8, 120.3, 119.0, 118.8, 100.8, 72.93, 72.86. HRMS *m/z* [M+H]^+^ calcd for C_17_H_14_N_2_OBr: 341.0289, found 341.0286, LC *t*_R_ = 3.28 min, > 98 % Purity.

***N*-(Benzo[*d*][1,3]dioxol-4-yl)-6-bromoquinolin-4-amine (27)** was obtained as a yellow solid (157 mg, 0.458 mmol, 74%). MP >270 °C decomp.; ^1^H NMR (400 MHz, DMSO-*d*_6_) δ 11.21 (s, 1H), 9.24 (d, J = 1.9 Hz, 1H), 8.57 (d, J = 6.9 Hz, 1H), 8.25 – 8.02 (m, 2H), 7.16 – 6.81 (m, 3H), 6.65 (d, J = 6.9 Hz, 1H), 6.11 (s, 2H). 13C NMR (101 MHz, DMSO-*d*_6_) δ 153.1, 148.8, 142.9, 141.5, 137.2, 136.5, 126.3, 122.6, 122.5, 120.0, 119.5, 119.1, 118.4, 107.9, 101.84, 101.80. HRMS m/z [M+H]^+^ calcd for C_16_H_12_N_2_O_2_Br: 343.0082, found 343.0072, LC t_R_ = 3.43 min, > 98% Purity.

**6-bromo-*N*-(1,3-dihydro-2-benzofuran-4-yl)quinolin-4-amine (28)** was obtained as a yellow/mustard solid (141 mg, 0.414 mmol, 67 %). MP Decomp. >220 °C; ^1^H NMR (400 MHz, DMSO-*d*_6_) δ 10.81 (s, 1H), 9.16 (d, *J* = 1.9 Hz, 1H), 8.49 (d, *J* = 6.6 Hz, 1H), 8.13 – 7.97 (m, 2H), 7.47 (t, *J* = 7.6 Hz, 1H), 7.35 (ddd, *J* = 21.1, 7.6, 0.9 Hz, 2H), 6.47 (d, *J* = 6.5 Hz, 1H), 5.10 (d, *J* = 2.0 Hz, 2H), 4.95 (d, *J* = 2.0 Hz, 2H). ^13^C NMR (100 MHz, DMSO-*d*_6_) δ 152.3, 144.9, 141.8, 139.6, 135.6, 135.4, 131.6, 129.3, 126.2, 124.7, 124.4, 120.2, 119.4, 119.1, 101.0, 73.0, 71.5. HRMS *m/z* [M+H]^+^ calcd for C_17_H_14_N_2_OBr: 341.0289, found 341.0281, LC *t*_R_ = 3.25 min, > 98 % Purity.

**6-bromo-*N*-(3-methoxyphenyl)quinazolin-4-amine (29)** was obtained as a light yellow solid (175 mg, 0.530 mmol, 86 %). MP 239-241 °C; ^1^H NMR (400 MHz, DMSO-*d*_6_) δ 11.79 (s, 1H), 9.34 (d, *J* = 2.0 Hz, 1H), 8.95 (s, 1H), 8.24 (dd, *J* = 8.9, 2.0 Hz, 1H), 7.98 (d, *J* = 8.9 Hz, 1H), 7.55 – 7.18 (m, 3H), 6.97 – 6.83 (m, 1H), 3.79 (s, 3H). ^13^C NMR (100 MHz, DMSO-*d*_6_) δ 159.4, 158.8, 151.1, 138.8, 138.1, 137.6, 129.5, 127.4, 122.0, 121.0, 116.7, 115.1, 112.0, 110.6, 55.3. HRMS *m/z* [M+H]^+^ calcd for C_15_H_13_N_3_OBr: 330.0242, found 330.0246, LC *t*_R_ = 4.80 min, > 98 % Purity.

**3-[(6-bromoquinazolin-4-yl)amino]phenol (30)** was obtained as a yellow solid (154 mg, 0.487 mmol, 79 %). MP Decomp. >250 °C; ^1^H NMR (400 MHz, DMSO-*d*_6_) δ 11.63 (s, 1H), 9.82 (s, 1H), 9.27 (d, *J* = 2.0 Hz, 1H), 8.94 (s, 1H), 8.23 (dd, *J* = 8.9, 2.0 Hz, 1H), 7.96 (d, *J* = 8.9 Hz, 1H), 7.43 – 6.92 (m, 3H), 6.76 (ddd, *J* = 8.1, 2.4, 1.0 Hz, 1H). ^13^C NMR (100 MHz, DMSO-*d*_6_) δ 158.7, 157.7, 151.1, 138.7, 138.1, 137.4, 129.4, 127.3, 122.1, 120.9, 115.2, 115.1, 113.9, 111.7. HRMS *m/z* [M+H]^+^ calcd for C_14_H_11_N_3_OBr: 316.0085, found 316.0081, LC *t*_R_ = 4.20 min, > 98 % Purity.

**6-bromo-*N*-[3-(2*H*-1,2,3,4-tetrazol-5-yl)phenyl]quinazolin-4-amine (31)** was obtained as a light beige solid (184 mg, 0.499 mmol, 81 %). ^1^H NMR (400 MHz, DMSO-*d*_6_) δ 11.21 (s, 1H), 9.18 (d, *J* = 2.0 Hz, 1H), 8.58 (d, *J* = 6.9 Hz, 1H), 8.23 (t, *J* = 1.9 Hz, 1H), 8.19 (dd, *J* = 9.0, 2.0 Hz, 1H), 8.13 (dt, *J* = 7.8, 1.3 Hz, 1H), 8.08 (d, *J* = 9.0 Hz, 1H), 7.80 (t, *J* = 7.9 Hz, 1H), 7.72 (ddd, *J* = 8.1, 2.2, 1.1 Hz, 1H), 7.02 (d, *J* = 6.9 Hz, 1H). ^13^C NMR (100 MHz, DMSO-*d*_6_) δ 153.8, 143.4, 138.1, 137.4, 136.7, 131.1, 127.6, 126.2, 126.0, 125.7, 123.4, 122.6, 120.1, 118.8, 100.8. MP Decomp. 300 °C; HRMS *m/z* [M+H]^+^ calcd for C_15_H_16_N_6_Br: 359.0620, found 369.0276, LC *t*_R_ = 3.10 min, > 98 % Purity.

**6-bromo-*N*-(3-(pentafluoro-λ^6^-sulfaneyl)phenyl)quinolin-4-amine (32)** was obtained as a colourless solid (200 mg, 0.470 mmol, 76 %). MP 290-292 °C; ^1^H NMR (400 MHz, DMSO-*d*_6_) δ 11.31 (s, 1H), 9.18 (d, *J* = 2.0 Hz, 1H), 8.61 (d, *J* = 6.9 Hz, 1H), 8.20 (dd, *J* = 9.0, 2.0 Hz, 1H), 8.12 (d, *J* = 9.0 Hz, 1H), 8.07 (t, *J* = 2.0 Hz, 1H), 7.95 (ddd, *J* = 8.0, 2.3, 1.2 Hz, 1H), 7.86 (dt, *J* = 8.1, 1.5 Hz, 1H), 7.81 (t, *J* = 8.0 Hz, 1H), 6.96 (d, *J* = 6.9 Hz, 1H). 13C NMR (100 MHz, DMSO-*d*_6_) δ 153.8, 153.5 (t, *J* = 16.7 Hz), 143.5, 138.2, 137.4, 136.7, 131.0, 129.0, 126.3, 124.3, 122.6, 122.6, 120.1, 118.9, 100.9. HRMS *m/z* [M+H]^+^ calcd for C_15_H_11_N_2_SF_5_Br: 424.9746, found 424.9736, LC *t*_R_ = 4.64 min, > 98 % Purity.

**6-bromo-*N*-[3-(*tert*-butoxy)phenyl]quinolin-4-amine (33)** was obtained as a colourless solid (170 mg, 0.458 mmol, 74 %). MP 280-282 °C; ^1^H NMR (400 MHz, DMSO-*d*_6_) δ 11.10 (s, 1H), 9.17 (d, *J* = 2.0 Hz, 1H), 8.54 (d, *J* = 6.9 Hz, 1H), 8.17 (dd, *J* = 9.0, 2.0 Hz, 1H), 8.08 (d, *J* = 9.0 Hz, 1H), 7.47 (t, *J* = 8.0 Hz, 1H), 7.19 (ddd, *J* = 7.9, 2.1, 0.9 Hz, 1H), 7.09 (t, *J* = 2.2 Hz, 1H), 7.03 (ddd, *J* = 8.2, 2.4, 0.9 Hz, 1H), 6.84 (d, *J* = 6.9 Hz, 1H), 1.35 (s, 9H). 13C NMR (101 MHz, DMSO-*d*_6_) δ 156.3, 154.0, 143.0, 137.6, 137.3, 136.6, 130.2, 126.2, 122.5, 122.4, 120.1, 119.8, 119.7, 118.6, 100.5, 78.8, 28.5 (s, 3C). HRMS *m/z* [M+H]^+^ calcd for C_19_H_20_N_2_OBr: 371.0759, found 371.0750, LC *t*_R_ = 4.54 min, > 98 % Purity.

**6-bromo-*N*-(3-*tert*-butylphenyl)quinolin-4-amine (34)** was obtained as a yellow solid (171 mg, 0.483 mmol, 78 %). MP 285-287 °C; ^1^H NMR (400 MHz, DMSO-*d*_6_) δ 11.08 (s, 1H), 9.17 (d, *J* = 2.0 Hz, 1H), 8.51 (d, *J* = 7.0 Hz, 1H), 8.17 (dd, *J* = 9.0, 2.0 Hz, 1H), 8.08 (d, *J* = 9.0 Hz, 1H), 7.86 – 7.37 (m, 3H), 7.31 (ddd, *J* = 7.4, 2.2, 1.3 Hz, 1H), 6.83 (d, *J* = 7.0 Hz, 1H), 1.32 (s, 9H). ^13^C NMR (100 MHz, DMSO-*d*_6_) δ 154.0, 152.9, 143.0, 137.3, 136.7, 136.5, 129.6, 126.2, 124.4, 122.4, 122.2, 119.77, 118.6, 100.3, 34.7, 31.0. HRMS *m/z* [M+H]^+^ calcd for C_19_H_20_N_2_Br: 355.0810, found 355.0800, LC *t*_R_ = 4.99 min, > 98 % Purity.

## Acknowledgements

The SGC is a registered charity (number 1097737) that receives funds from AbbVie, Bayer Pharma AG, Boehringer Ingelheim, Canada Foundation for Innovation, Eshelman Institute for Innovation, Genome Canada, Innovative Medi-cines Initiative (EU/EFPIA) [ULTRA-DD grant no. 115766], Janssen, Merck KGaA Darmstadt Germany, MSD, Novartis Pharma AG, Ontario Ministry of Economic Development and Innovation, Pfizer, São Paulo Research Foundation-FAPESP, Takeda, and Wellcome [106169/ZZ14/Z]. We also thank CSC - IT Center for Science Ltd. Finland for the use of their facilities, software licenses, computational resources and the Biocenter Finland/DDCB for financial support. The authors thank Prof. Lee Graves (University of North Carolina at Chapel Hill) for useful discussions. We are grateful to Dr. Brandie Ehrmann for LC-MS/HRMS support provided by the Mass Spectrometry Core Laboratory at the University of North Carolina at Chapel Hill. We also thank the EPSRC UK National Crystallography Service for funding and collection of the crystallographic data for **13, 14, 25, 27** and **29**

## Entry for the Table of Contents

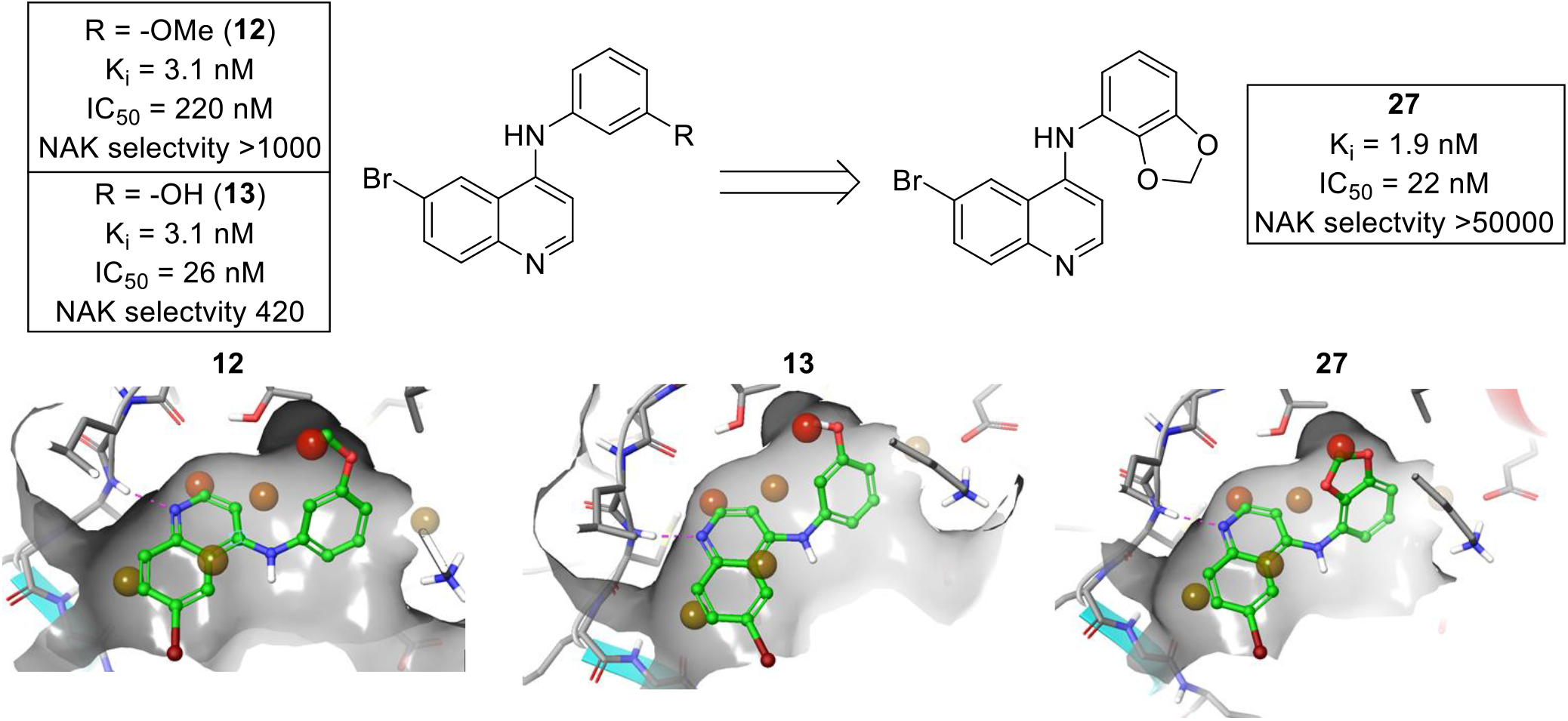

Water networks within kinase inhibitor design and drug discovery more widely are generally poorly understood. Here we elucidate the properties of a proposed water network with cyclin G associated kinase (GAK). A targeted library of point mutations was used to build a body of evidence to support the prospective targeting of the GAK water network. This can be used as not only a tool to more potent GAK inhibitors but also enhance wider inhibitor design.

## References

[1] E. D. Scheeff, P. E. Bourne, PLoS Comput Biol. 2005, 5, e49.

[2] Z. A. Knight, K. M. Shokat, Chemistry & Biology 2005, 12, 621–637.

[3] G. Manning, D. B. Whyte, R. Martinez, T. Hunter, S. Sudarsanam, Science 2002, 298, 1912–1934.

[4] P. Badrinarayan, G. N. Sastry, Curr Pharm Des. 2013, 19, 4714–4738.

[5] S. Lu, X. He, D. Ni, J. Zhang J. Med. Chem. 2019, 62, 6405–6421.

[6] Q. Liu, Y. Sabnis, Z. Zhao, T. Zhang, S. J. Buhrlage, L. H. Jones, N. S. Gray Chem Biol. 2013, 20, 146–159.

[7] W. D. Jang, J. Kim, N. S. Kang, Journal of Molecular Liquids 2014, 191, 37–41.

[8] W. D. Jang, M. H. Lee, N. S. Kang, Journal of Molecular Liquids 2016, 221, 361–322.

[9] F. Heider, T. Pantsar, M. Kudolo, F. Ansideri, A. De Simone, L. Pruccoli, T. Schneider, M. I. Goettert, A. Tarozzi, V. Andrisano, S. A. Laufer, P. Koch ACS Med. Chem. Lett. 2019, 10, 1407–1414

[10] S. Riniker, L. J. Barandun, F. Diederich, O. Krämer, A. Steffen, W. F. van Gunsteren, J Comput Aided Mol Des. 2012, 26, 1293–1309.

[11] R. Abel, L. Wang, R. A. Friesner, B. J. Berne, J Chem Theory Comput. 2010, 6, 2924–2934.

[12] Y. Yang, F. C. Lightstone, S. E. Wong, Expert Opin Drug Discov. 2013, 8, 277–287.

[13] D. D. Robinson, W. Sherman, R. Farid, ChemMedChem. 2010, 5, 618–627.

[14] S. Kannan, M. R. Pradhan, G. Tiwari, W. C. Tan, B. Chowbay, E. H. Tan, D. S. Tan, C. Verma, Sci Rep. 2017, 7, 1540.

[15] R. Horbert, B. Pinchuk, E. Johannes, J. Schlosser, D. Schmidt, D. Cappel, F. Totzke, C. Schächtele, C. Peifer, J Med Chem. 2015, 58, 170–182.

[16] N. M. Levinson, S. G. Boxer, Nat Chem Biol. 2014, 10, 127–132.

[17] C. R. M. Asquith, T. Laitinen, J. M. Bennett, P. H. Godoi, M. P. East, G. J. Tizzard, L. M. Graves, G. L. Johnson, R. E. Dornsife, C. I. Wells, J. M. Elkins, T. M. Willson, W. J. Zuercher, ChemMedChem 2018, 13, 48–66.

[18] C. R. M. Asquith, B. T. Berger, J. Wan, J. M. Bennett, S. J. Capuzzi, D. J. Crona, D. H. Drewry, M. P. East, J. M. Elkins, O. Fedorov, P. H. Godoi, D. M. Hunter, S. Knapp, S. Müller, C. D. Torrice, C. I. Wells, H. S. Earp, T. M. Willson, W. J. Zuercher, J. Med. Chem. 2019, 62, 2830–2836.

[19] C. R. M. Asquith, D. K. Treiber, W. J. Zuercher, Bioorg. Med. Chem. Lett. 2019, 29, 1727–1731.

[20] C. R. M. Asquith, T. Laitinen, J. M. Bennett, C. I. Wells, J. M. Elkins, W. J. Zuercher, G. J. Tizzard, A. Poso, ChemMedChem. 2020, 15, 26–49.

[21] F. J. Sorrell, M. Szklarz, K. R. Abdul Azeez, J. M. Elkins, S. Knapp, Structure 2016, 24, 401–411.

[22] H. Shimizu, I. Nagamori, N. Yabuta, H. Nojima J Cell Sci. 2009, 122, 3145–3152.

[23] N. Dzamko, J. Zhou, Y. Huang, G. M. Halliday, Front. Mol. Neurosci. 2014, 7, 57.

[24] M. Susa, E. Choy, X. Liu, J. Schwab, F. J. Hornicek, H. Mankin, Z. Duan, Mol. Cancer Ther. 2010, 9, 3342–3350.

[25] M. R. Ray, L. A. Wafa, H. Cheng, R. Snoek, L. Fazli, M. Gleave, P. S. Rennie, Int. J. Cancer 2006, 118, 1108–1119.

[26] S. Knapp, P. Arruda, J. Blagg, S. Burley, D. H. Drewry, A. Edwards, D. Fabbro, P. Gillespie, N. S. Gray, B. Kuster, K. E. Lackey, P. Mazzafera, N. C. Tomkinson, T. M. Willson, P. Workman, W. J. Zuercher, Nat. Chem. Biol. 2013, 9, 3–6.

[27] S. Kovackova, L. Chang, E. Bekerman, G. Neveu, R. Barouch-Bentov, A. Chaikuad, C. Heroven, M. Sala, S. De Jonghe, S. Knapp, S. Einav, P. Herdewijn, J. Med. Chem. 2015, 58, 3393–3410.

[28] S. Y. Pu, R. Wouters, S. Schor, J. Rozenski, R. Barouch-Bentov, L. I. Prugar, C. M. O’Brien, J. M. Brannan, J. M. Dye, P. Herdewijn, S. De Jonghe, S. Einav, J Med Chem. 2018, 61, 6178–6192.

[29] C. H. Arrowsmith, J.E. Audia, C. Austin, J. Baell, J. Bennett, J. Blagg, C. Bountra, P. E. Brennan, P. J. Brown, M. E. Bunnage, C. Buser-Doepner, R. M. Campbell, A. J. Carter, P. Cohen, R. A. Copeland, B. Cravatt, J. L. Dahlin, D. Dhanak, A. M. Edwards, M. Frederiksen, S. V. Frye, N. Gray, C. E. Grimshaw, D. Hepworth, T. Howe, K. V. Huber, J. Jin, S. Knapp, J. D. Kotz, R. G. Kruger, D. Lowe, M. M. Mader, B. Marsden, A. Mueller-Fahrnow, S. Müller, R. C. O’Hagen, J. P. Overington, D. R. Owen, S. H. Rosen-berg, B. Roth, R. Ross, M. Schapira, S. L. Schreiber, B. Shoicet, M. Sundstrçm, G. Superti-Furga, J. Taunton, L. Toledo-Sherman, C. Walpole, M. A. Walters, T. M. Wilson, P. Workman, R. N. Young, W. J. Zuercher, Nat. Chem. Biol. 2015, 11, 536–542.

[30] M. A. Fabian, W. H. Biggs III, D. K. Treiber, C. E. Atteridge, M.D. Azimioara, M. G. Benedetti, T. A. Carter, P. Ciceri, P. T. Edeen, M. Floyd, J. M. Ford, M. Galvin, J. L. Gerlach, R. M. Grotzfeld, S. Herrgard, D. E. Insko, M. A. Insko, A. G. Lai, J. M. Lelias, S. A. Mehta, Z. V. Milanov, A. M. Velasco, L. M. Wodicka, H. K. Patel, P. P. Zarrinkar, D. J. Lockhart, Nat. Biotechnol. 2005, 23, 329–336.

[31] R. Jr. Roskoski, Pharmacol Res. 2016, 103, 26–48.

[32] R. Newton, K. A. Bowler, E. M. Burns, P. J. Chapman, E. E. Fairweather, S. J. R. Fritzl, K. M. Goldberg, N. M. Hamilton, S.V. Holt, G. V. Hopkins, S. D. Jones, A. M. Jordan, A. J. Lyons, H. Nikki March, N. Q. McDonald, L. A. Maguire, D. P. Mould, A.G. Purkiss, H. F. Small, A. I. J. Stowell, G. J. Thomson, I. D. Waddell, B. Waszkowycz, A. J. Watson, D. J. Ogilvie, Eur J Med Chem. 2016, 13, 20–32.

[33] C. R. M. Asquith, N. Fleck, C. D. Torrice, D. J. Crona, C. Grundner, W. J. Zuercher, Bioorg. Med. Chem. Lett. 2019, 18, 2695–2699.

[34] C. R. M. Asquith, K. M. Naegeli, M. P. East, T. Laitinen, T. M. Havener, C. I. Wells, G. L. Johnson, D. H. Drewry, W. J. Zuercher, D. C. Morris, J Med Chem. 2019, 62, 4772–4778.

[35] T. Machleidt, C. C. Woodroofe, M. K. Schwinn, J. Méndez, M. Robers, K. Zimmerman, P. Otto, D. L. Daniels, T. A. Kirkland, K. V. Wood, ACS Chem Biol. 2015, 10, 1797–1804.

[36] J. D. Vasta, C. R, Corona, J. Wilkinson, C. A. Zimprich, J. R. Hartnett, M. R. Ingold, K. Zimmerman, T. Machleidt, T. A. Kirkland, K. G. Huwiler, R. F. Ohana, M. Slater, P. Otto, M. Cong, I. Wells, B. T. Berger, T. Hanke, C. Glas, K. Ding, D. H. Drewry, K. V. M. Huber, T. M. Willson, S. Knapp, S. Müller, P.L. Meisenheimer, F. Fan, K. V. Wood, M. B. Robers, Cell Chem. Biol. 2018, 25, 206–214.

[37] W. F. Baitinger, P. von R.Schleyer, T. S. S. R. Murty, L. Robinson, Tetrahedron 20, 1964, 1635–1647.

[38] L. Pauling, J. Am. Chem. Soc. 1932, 54, 3570–3582.

[39] P. R. Savoie, J. T. Welch, Chem Rev. 2015, 115, 1130–1190.

[40] Schrödinger Maestro software package (Small-Molecule Drug Discovery Suite 2018–4, Schrödinger, LLC, New York, NY, 2018).

[41] P. Politzer, J. S. Murray, T. Clark, Phys. Chem. Chem. Phys. 2013, 15, 11178–11189.

[42] L. Wang, B. J. Berne, R. A. Friesner, Proc Natl Acad Sci U S A. 2011, 108, 1326–1330.

[43] a) F. M. Ferguson, N. S. Gray, Nat Rev Drug Discov. 2018, 17, 353–377.; b) http://www.brimr.org/PKI/PKIs.htm

[44] P. Cohen, D. R. Alessi, ACS Chem. Biol. 2013, 8, 96–104.

[45] A. F. Rudolf, T. Skovgaard, S. Knapp, L. J. Jensen, J. Berthelsen, PLoS One 2014, 9, e98800.

[46] M. I. Davis, J. P. Hunt, S. Herrgard, P. Ciceri, L. M. Wodicka, G. Pallares, M. Hocker, D. K. Treiber, P. P. Zarrinkar, Nat. Biotechnol. 2011, 29, 1046–1051.

[47] H. Ma, S. Deacon, K. Horiuchi, Expert Opin. Drug Discovery 2008, 3, 607–621.

[48] a) A. J. Kooistra, G. K. Kanev, O. P. van Linden, R. Leurs, I. J. de Esch, C. de Graaf, Nucleic Acids Res. 2016, 44, D365–D371.; b) O. P. van Linden, A. J. Kooistra, R. Leurs, I. J. de Esch, C. de Graaf, J. Med. Chem. 2014, 57, 249–277.

[49] D. Robinson, T. Bertrand, J. C. Carry, F. Halley, A. Karlsson, M. Mathieu, H. Minoux, M. A. Perrin, B. Robert, L. Schio, W. Sherman, J Chem Inf Model. 2016, 56, 886–894.

[50] P. Czodrowski, G. Hölzemann, G. Barnickel, H. Greiner, D. Musil, J Med Chem. 2015, 58, 457–465.

[51] V. Myrianthopoulos, M. Kritsanida, N. Gaboriaud-Kolar, P. Magiatis, Y. Ferandin, E. Durieu, O. Lozach, D. Cappel, M. Soundararajan, P. Filippakopoulos, W. Sherman, S. Knapp, L. Meijer, E. Mikros, A. L. Skaltsounis, ACS Med Chem Lett. 2013, 4, 22–26.

[52] N. Ohbayashi, K. Murayama, M. Kato-Murayama, M. Kukimoto-Niino, T. Uejima, T. Matsuda, N. Ohsawa, S. Yokoyoma, H. Nojima, M. Shirouzu, ChemistryOpen. 2018, 7, 721–727.

[53] T. Young, R. Abel, B. Kim, B. J. Berne, R. A. Friesner, Proc Natl Acad Sci U S A. 2007, 104, 808–813.

[54] R. Abel, T. Young, R. Farid, B. J. Berne, R. A. Friesner, J. Am. Chem. Soc. 2008, 130, 2817–2831

[55] D. Cappel, W. Sherman, T. Beuming, Curr Top Med Chem. 2017, 17, 2586–2598.

[56] E. Harder, W. Damm, J. Maple, C. Wu, M. Reboul, J. Y. Xiang, L. Wang, D. Lupyan, M. K. Dahlgren, J. L. Knight, J. W. Kaus, D. S. Cerutti, G. Krilov, W. L. Jorgensen, R. Abel, R. A. Friesner, J Chem Theory Comput. 2016, 12, 281–296.

[57] C. R. M. Asquith, J. M. Bennett, L. Su, T. Laitinen, J. M. Elkins, J. E. Pickett, C. I. Wells, Z. Li, T. M. Willson, W. J. Zuercher, Molecules 2019, 24, 4016.

